# Thienothiazolocarboxamides: rapidly cidal agents causing catastrophic thiol collapse in *Mycobacterium tuberculosis*

**DOI:** 10.64898/2026.11.06.764870

**Authors:** Sangmi Oh, Gareth Prosser, Helena I. M. Boshoff, Simon R. Green, Neha Quadir, Jose Santinni O. Roma, Ashish Kumar, Nilesh Mukundan, Sarah Hanson, Jansy Sarathy, Kirsteen I. Buchanan, Peter C. Ray, Paul G. Wyatt, Thomas R. Ioerger, Adrian L. Schwan, Laura A. T. Cleghorn, Clifton E. Barry

**Author notes:** These authors contributed equally (Sangmi Oh, Gareth Prosser).

## Abstract

Thienothiazolocarboxamides (TTCs) were discovered as a hit in a whole cell screen for new antituberculosis agents. These molecules were rapidly cidal under both replicating and non-replicating conditions and killed *Mycobacterium tuberculosis (Mtb)* in macrophages. The heterobicyclic core of this scaffold was essential for activity and resistance to these agents was mediated by loss of EthA, a flavin containing monooxygenase also responsible for the bioactivation of ethionamide. EthA-mediated activation of TTCs in whole cells was found to produce mycothiol adducts and RNAseq profiles revealed a mechanism of action identical to the fungal mycotoxins patulin and nidulalin A. EthA oxidizes the sulfur of the thiazole ring to create a relatively stable sulfenic acid that reacts with cellular thiols. This unique, replication-independent, mechanism suggests the possibility that these molecules may contribute significantly to shortening the duration of tuberculosis chemotherapy.

---

Tuberculosis (TB) is the leading cause of death from a single infectious agent. In 2024, 10.7 million people fell ill with the disease and 1.23 million people died; concerningly, of the patients tested for resistance, ∼5% showed resistance to at least one frontline drug^1^. Recent reductions in global funding for TB control programs, particularly the dismantling of USAID, is projected to dramatically exacerbate the pace of this epidemic resulting in up to an additional 14 million deaths by 2030^2,3^.

The newest clinical agents for TB treatment such as bedaquiline, pretomanid and linezolid were all identified through traditional phenotypic screening against actively replicating organisms^4–6^. While these drugs are clinically approved, their use so far has been limited to treating patients resistant to the frontline therapies^7^. The development of shorter duration regimens is a major goal of current drug development activities in TB although predicting treatment duration from preclinical data remains a challenge^8^. Traditionally treatment shortening has been accomplished empirically by employing combinations of new agents in Phase 2B/3 clinical trials. Recently, shortening treatment to four-months has been successful utilizing a combination of rifapentine (a longer half-life analog of rifamycin) and moxifloxacin^9^.

CRISPRi analysis to rank target vulnerability has focussed efforts on highly sensitive targets that can be exploited in target-based screening approaches^10^ but these genetic tools have primarily been applied in the context of rapidly growing cells. The duration of treatment is thought to be driven by the activity of the agents in the regimen against non-replicating *Mtb* within caseous granulomas^11^. In this microenvironment *Mtb* becomes drug tolerant to many drugs making treatment-shortening an often-elusive goal.

In previous work, we sampled fungi from core anaerobic regions of sphagnum bogs to understand what strategies organisms used to compete with *Mtb* under similar growth-restrictive conditions^12^. When co-cultured with *Mtb,* five different fungi upregulated biosynthetic genes that corresponded to three structurally different mycotoxins that all killed *Mtb* by causing catastrophic thiol collapse. In the acidic, hypoxic environment of a sphagnum peat core selection of this mechanism by different fungi in response to the presence of *Mtb* suggests that this may be a particularly sensitive aspect of metabolism under these conditions which are similar to the conditions observed in caseum.

## Results

### TTCs have potent antitubercular activity

As part of our effort to identify starting points for new drugs, a phenotypic screening program was performed, using a 530,000-compound library, provided by Medicines for Malaria Venture, that identified the potent singleton **1** (Fig. 1a). Medicinal chemistry optimization yielded a more soluble candidate **2** (**DDU384**, reference to companion paper)^13^. During this effort, this scaffold was independently identified, and a separate lead compound (**42**) was identified (Fig. 1a)^14^. Both lead compounds showed intramacrophage activity equal to or better than their *in vitro* MIC.

**Fig. 1:**
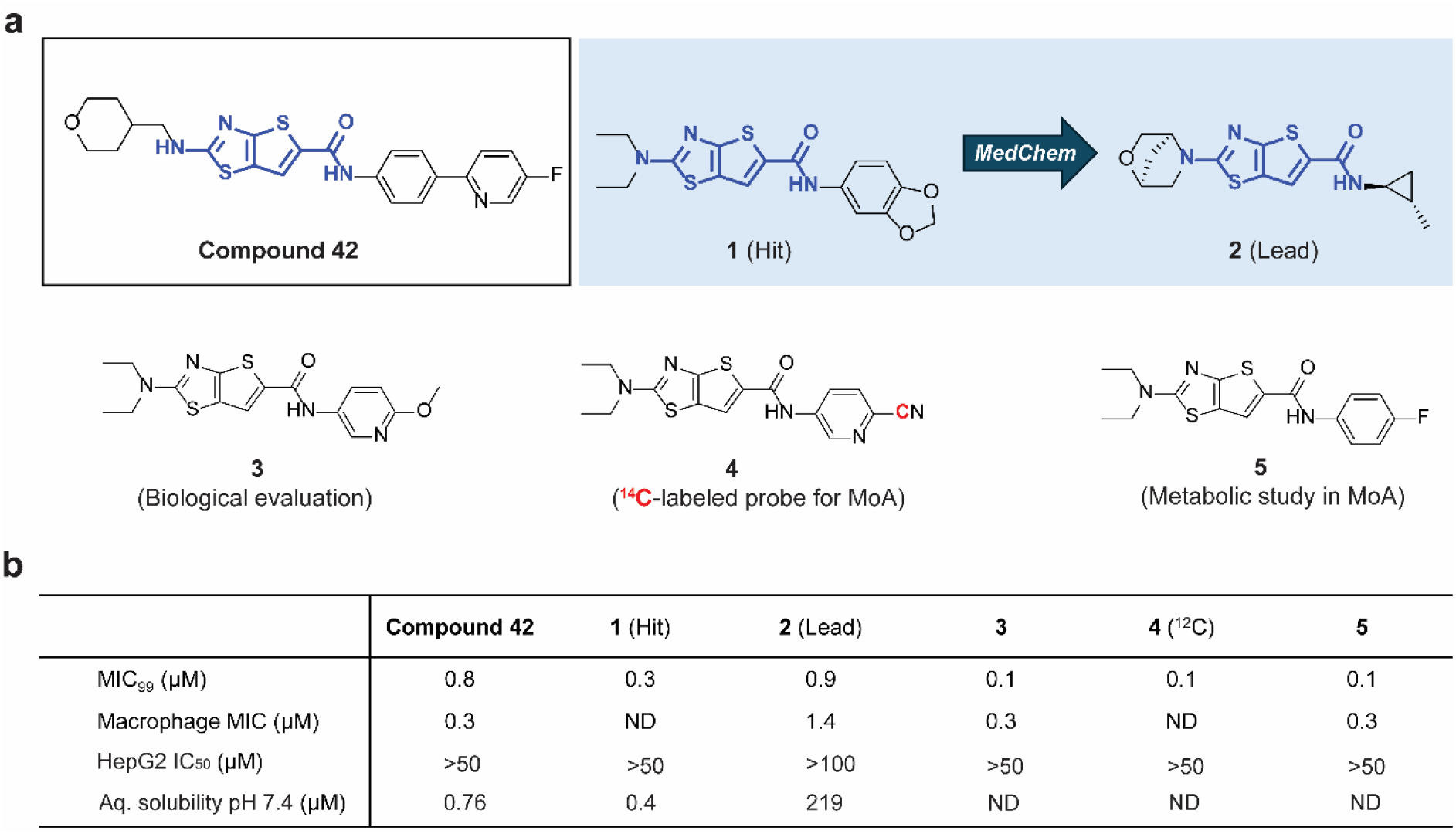
Summary of key TTC analogs. **a,** Chemical structure of TTC analogs investigated in this study. **b,** MIC is the minimum concentration required to inhibit the growth of *Mtb* (H37Rv) in liquid culture by 99% compared to untreated control; Macrophage MIC is the concentration required to inhibit 90% of the luminescent signal from a luciferase-expressing *Mtb* strain growing in THP-1 monocytes; HepG2 inhibitory concentration (IC50) is the concentration required to inhibit growth of HepG2 cells by 50%; Aqueous (Aq.) solubility at pH 7.4.

### Microbiological evaluation of antitubercular activity

The lead compound **2** was bactericidal causing a four to five log decrease in colony forming units (CFU) in a standard bactericidal activity assay (Fig. 2a). Compound **2** also showed almost four logs of killing in a starvation-induced non-replicating cidality assay (Fig. 2b)^15,16^. Compound **2** was cidal to non-replicating TB in the very difficult to treat *ex vivo* rabbit caseum model (Fig. 2c)^17^. In addition, compound **3** was comparable to rifampicin against bacteria within infected macrophages (Fig. 2d).

**Fig. 2:**
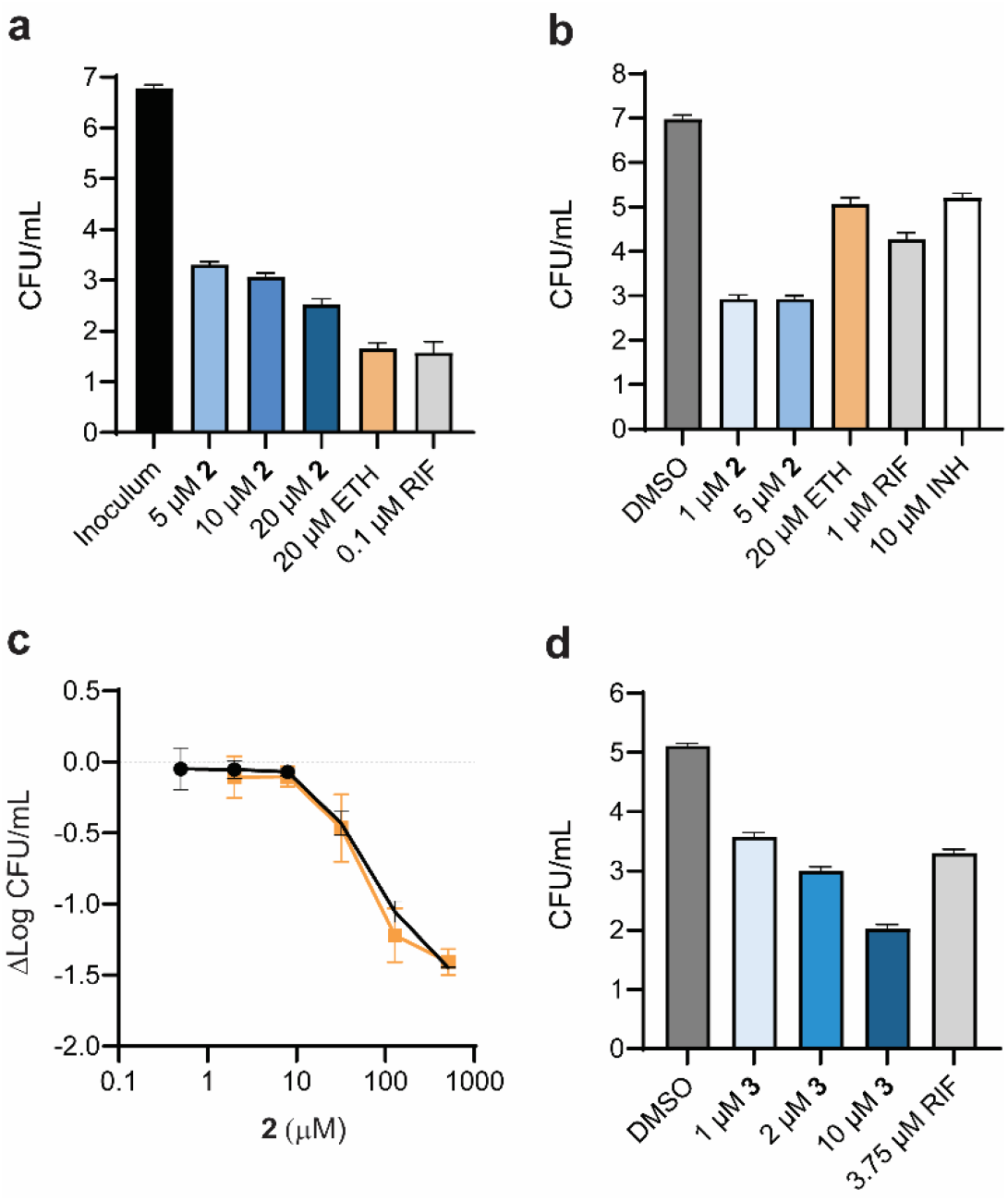
Microbiological characterization of 2. **a,** Minimum bactericidal activity following 7 days of exposure. **b,** Loebel (non-replicating) killing assay. **c,** Killing in *ex vivo* rabbit caseum, means and standard deviations are shown for three technical replicates from two independent experiments. **d,** Killing in infected macrophages, mean and standard deviations are show for three individual wells.

### TTCs are incorporated into stable cellular metabolites after EthA activation

Using a reporter assay^18,19^ TTCs were shown to not inhibit cell wall biosynthesis (Fig. 3a). *In vitro* selected resistant mutants showed predominantly loss of function mutations in the *ethA* gene, which was confirmed by evaluating 48 resistant mutants, of which 47 had mutations in *ethA* and one had a mutation in its regulator *ethR* (Fig. 3b). Overexpression of wild type EthA within resistant strains, fully restored sensitivity to the series (Fig. 3c) confirming its role in mechanism of resistance. Other antitubercular drugs are activated by EthA including ethionamide^20^, isoxyl^21^ and thiacetazone^22^ as well as a variety of non-clinical thiol-containing hits^23,24^. This resistance mechanism suggested that TTCs are prodrugs which are activated to form reactive metabolites in cells with lethal consequences. Ethionamide, for example, is activated to form an intracellular metabolite that accumulates within cells and inhibits mycolic acid biosynthesis^20,25^. The TTCs differed from ethionamide in not triggering a cell wall damage response, having activity in the non-replicating Loebel assay, and having activity in caseum suggesting the target was cell growth independent.

**Fig. 3:**
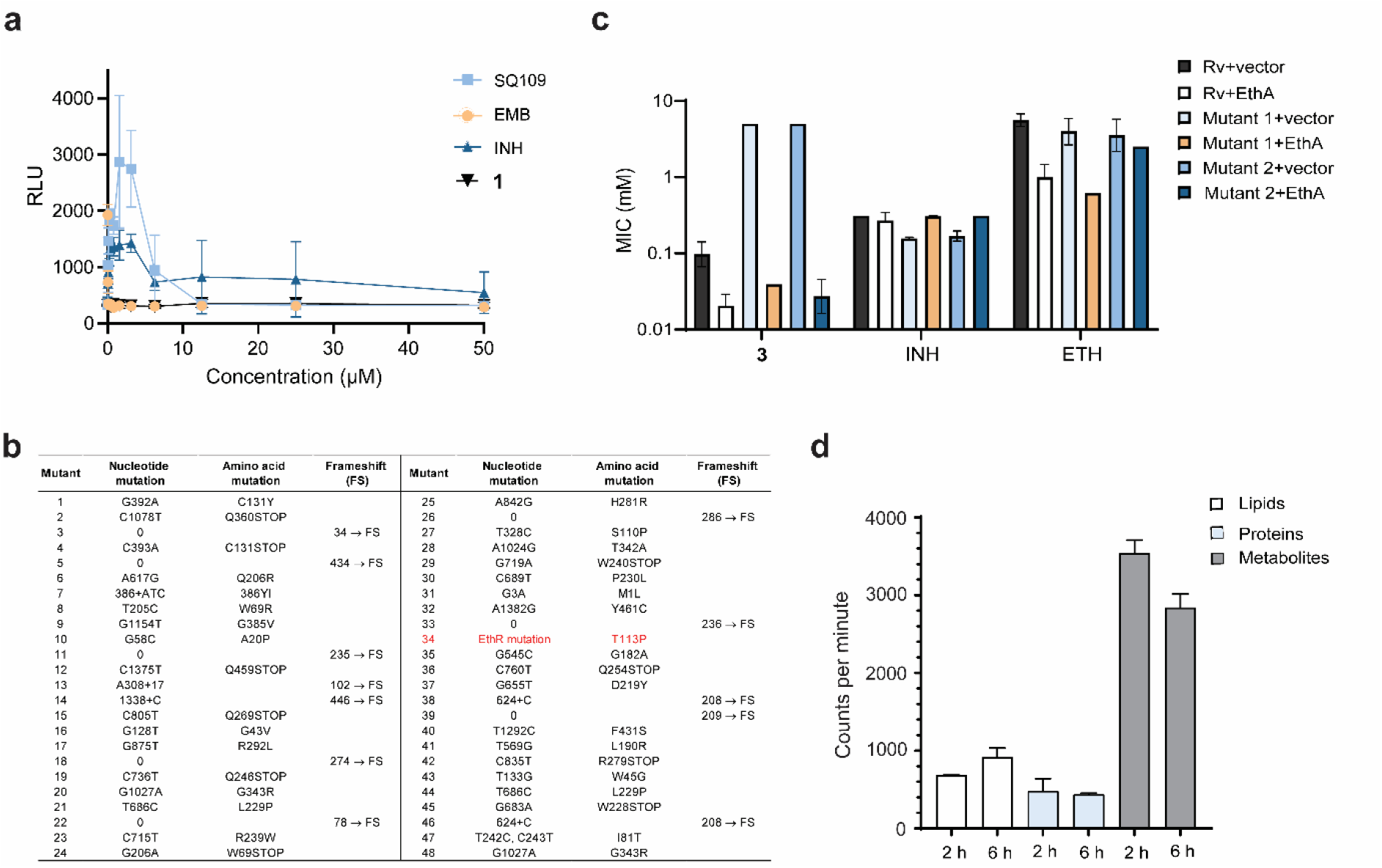
TTC mechanism of action studies. **a,** TTCs do not inhibit cell wall synthesis using a cell wall sensitive promoter fusion. **b,** Resistance to TTCs is mediated by loss of function mutations in EthA. Forty-seven independent mutants show mutation in EthA and one in EthR (T113P). **c,** Inserting a wild type copy of EthA back into two separate strains expressing mutant EthAs restores sensitivity to TTCs. **d,** A ^14^C-labeled TTC analog **4** shows that activation primarily results in incorporation into small molecule metabolites in *Mtb* cells.

We synthesized a bifunctional probe **6** based on an active analog of this series with an aryl azide and an alkyne handle (Supplementary Fig. 1). Labelling of whole cells of *Mtb* with this probe followed by UV irradiation and copper catalysed click chemistry revealed dozens of labelled targets. This labelling was increased with EthA overexpression and abolished in an EthA mutant resistant to TTCs. The labelling pattern of this probe was distinct from that obtained with Coomassie staining suggesting only a subset of proteins were being labelled. To establish the nature of potential targets labelled by TTCs, we also synthesized a ^14^C- labeled TTC (**4**, Fig. 1a). Following treatment of whole cells of *Mtb* with **4** we lysed cells and performed a chloroform:methanol:water extraction to separate lipids, protein and free metabolite fractions. Scintillation counting showed that the majority of the radioactivity mapped to the metabolite fraction (Fig. 3d, Supplementary Fig. 2). The labelled metabolite fraction could be ascribed to TTCs end products after EthA-mediated oxidation or the products of reaction with cellular metabolites. Thin-layer chromatography analysis of the radiolabelled metabolite fraction indicated that the ^14^C-labeled metabolites did not include the parent compound suggesting the formation of distinct metabolites.

### Chemical mechanism of EthA mediated activation of TTCs

EthA is a Baeyer-Villiger monooxygenase that utilizes a flavin to activate molecular oxygen to the flavin hydroperoxide^26–28^. To model this activation, we used an organic peroxide, *m*-chloroperoxybenzoic acid (mCPBA) to oxidize **5** in the presence of excess ethanethiol to trap reactive intermediates. We were not able to unambiguously determine the structure from ^1^H-NMR and therefore turned to single molecule X-ray crystallography to determine the structure (Supplementary Fig. 3). This revealed initial oxidation of the sulfur in the thiazole ring, presumably because it is slightly more nucleophilic than the sulfur in the thiophene, which was subsequently trapped with two equivalents of ethanethiol.

We noted that *Mtb* cells treated with various TTC analogs turned bright yellow with a strong absorbance at 420 nm. Extraction of TTC treated cultures and LC-MS analysis revealed two predominant molecular ions with a strong absorbance at 420 nm, M+191 (**M191**) and M+62 (**M62**, M is the parent molecular weight). Large cultures of *Mtb* followed by isolation of the two yellow metabolites provided purified materials that were again difficult to unambiguously characterize by NMR. Single crystal X-ray analysis revealed the structures as shown in Fig. 4a. Surprisingly the product structure suggested initial oxidation of the thiophene ring, not the thiazole observed in chemical oxidation. **M191** was the major product and contains a *N*-acetyl cysteine moiety in the structure, which is the key amino acid found in mycothiol, and two additional methyl groups. Mycothiol is the mycobacterial analog of the eukaryotic glutathione and serves to detoxify alkylating agents, reactive oxygen species and maintain the strongly reducing environment inside the cytoplasm^29^. Mycothiol reacts with alkylating agents and is then cleaved by mycothiol conjugate amidase to release the *N*-acetyl cysteine adduct for secretion^30^.

**Fig. 4:**
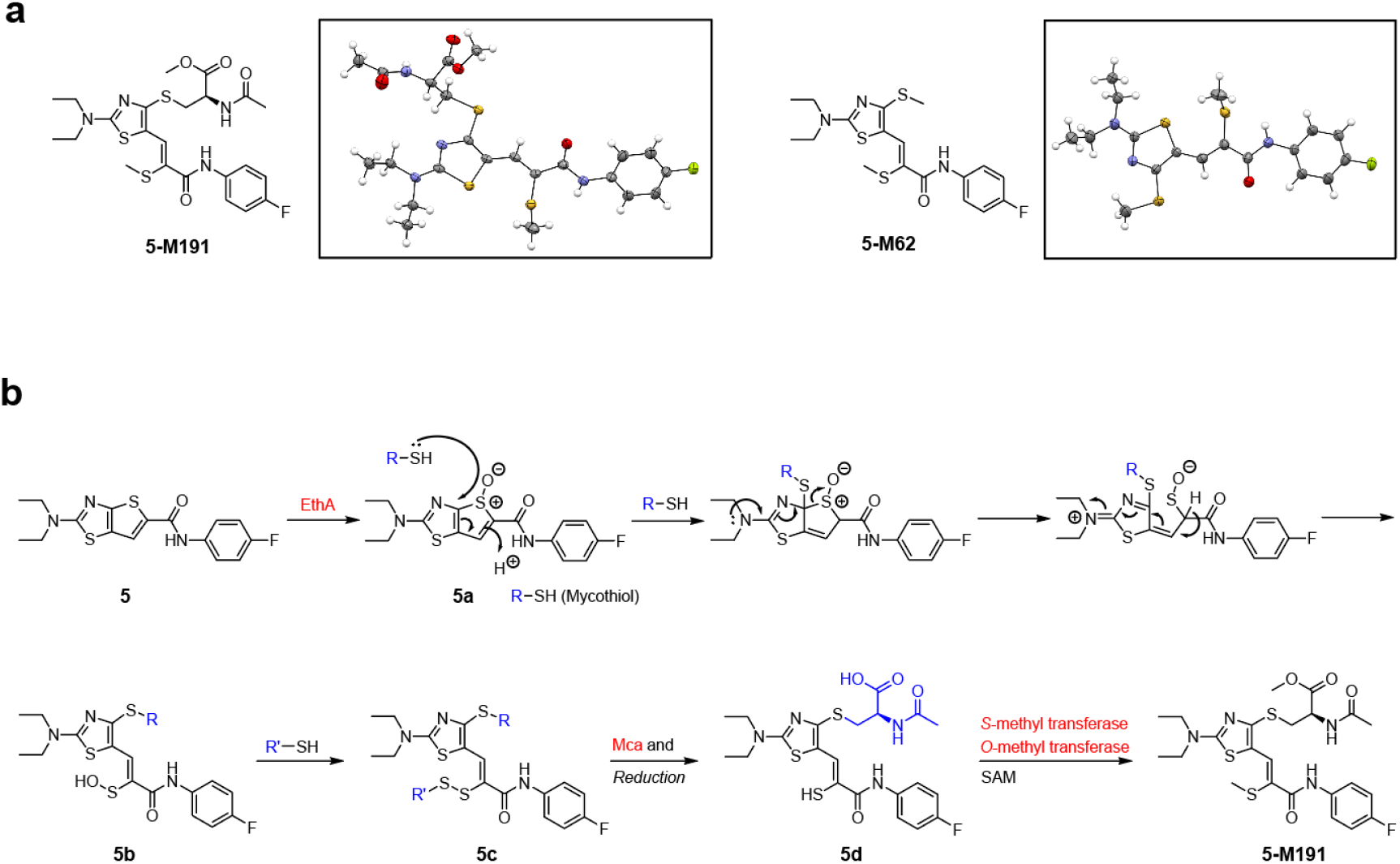
Identification of metabolites. **a,** Confirmed chemical structures of two major metabolites, **M191** and **M62** (**5-M191** and **5-M62** derived from **5**). **b,** Proposed metabolic degradation of EthA-activated TTCs by thiol quenching inside cells.

Mycothiol is biosynthesized in four steps starting with the glycosyltransferase MshA which links inositol-1-phosphate to *N*-acetylglucosamine. To demonstrate that activated TTCs react with mycothiol, we created an *Mtb* strain in which *mshA* was deleted. Deletion of *mshA* abolishes mycothiol biosynthesis^30^. When incubated with TTCs, this strain no longer produced the **M191** product peak demonstrating that this product requires the presence of mycothiol (Supplementary Fig. 4). Ergothioneine is the other major low molecular weight thiol present in *Mtb* and is biosynthesized from histidine by methylation to form hercynine by the enzyme EgtD. To understand the role of ergothioneine in the observed metabolites we generated a deletion in *egtD* and found that the level of **M191** and **M62** remained virtually unchanged (Supplementary Fig. 4). The origin of the two methyl groups in the product was established by labelling *Mtb* cells with CD_3_-methionine followed by treatment with a TTC analog. Analysis of the reaction product revealed M+3 and +6 peaks supporting incorporation of two *S*-adenosyl-L-methionine derived methyl groups into the final metabolites, **M191** and **M62** (Supplementary Fig. 5).

### Mechanism of enzymatic activation of TTCs by EthA

When we incubated active analogs with purified EthA protein in the presence of synthetic methylated *N*-acetyl cysteine (Me-NAC), we observed an initial oxidation product with one equivalent of Me-NAC by LC-MS, which was predicted to be the sulfenic acid **I1** (Supplementary Fig. 6, **5-I1**). We also observed a second lower abundance product with the molecular mass which we assigned as disulfide **I2**. Surprisingly sulfenic acid **5-I1** was stable enough to isolate by preparative TLC allowing purification of the sulfenic acid which was then reacted with dimedone and characterized. Dimedone reacts with sulfenic acids to form a stable covalent adduct^30,31^ although in the case of **5-I1** this reaction was slow suggesting an unusually stable sulfenic acid. To verify the other intermediate **5-I2**, we purified this compound and observed disulfide interchange to a dimeric form. Treatment of **5-I2** and its dimer with iodomethane in the presence of base generated the **5-M191** (**M191** derived from **5**) product, which is the major metabolite detected in cells.

Based on the results, we propose that initial oxidation of the thiophene by EthA in whole cells of *Mtb* is followed by nucleophilic attack by cellular thiols dominated by the major redox buffer mycothiol generating a relatively stable sulfenic acid (**5b**, Fig. 4b). This product is subsequently alkylated by a second cellular thiol including cysteine-containing proteins some of which are essential. The resulting adduct is then a substrate for mycothiol conjugate amidase (Mca) that cleaves the disaccharide from mycothiol leaving the *N*-acetyl cysteine. Two methyl transferases then use SAM-derived methyl groups to provide the final **5-M191** product.

### Mechanism of cell killing; collapse of thiol homeostasis

To further probe the mechanism of killing, we performed RNAseq analysis on *Mtb* cultures treated with TTCs and ethionamide. The transcriptional response of *Mtb* to treatment with **2** was dominated by members of the SigH regulon. The SigH regulon plays a key role in the response to oxidative stress in *Mtb*^32^ and includes SigE as well as the thioredoxin reductase TrxB2, ClpB involved in proteostasis, the mycothiol-dependent reductase Mrx2^33^, and the sulfate ABC transport system SubI/CysTWA^34^. Previous studies have characterized the SigH regulon as encompassing 48 genes^35^ of which 36 were upregulated by **2** (Fig. 5a).

**Fig. 5:**
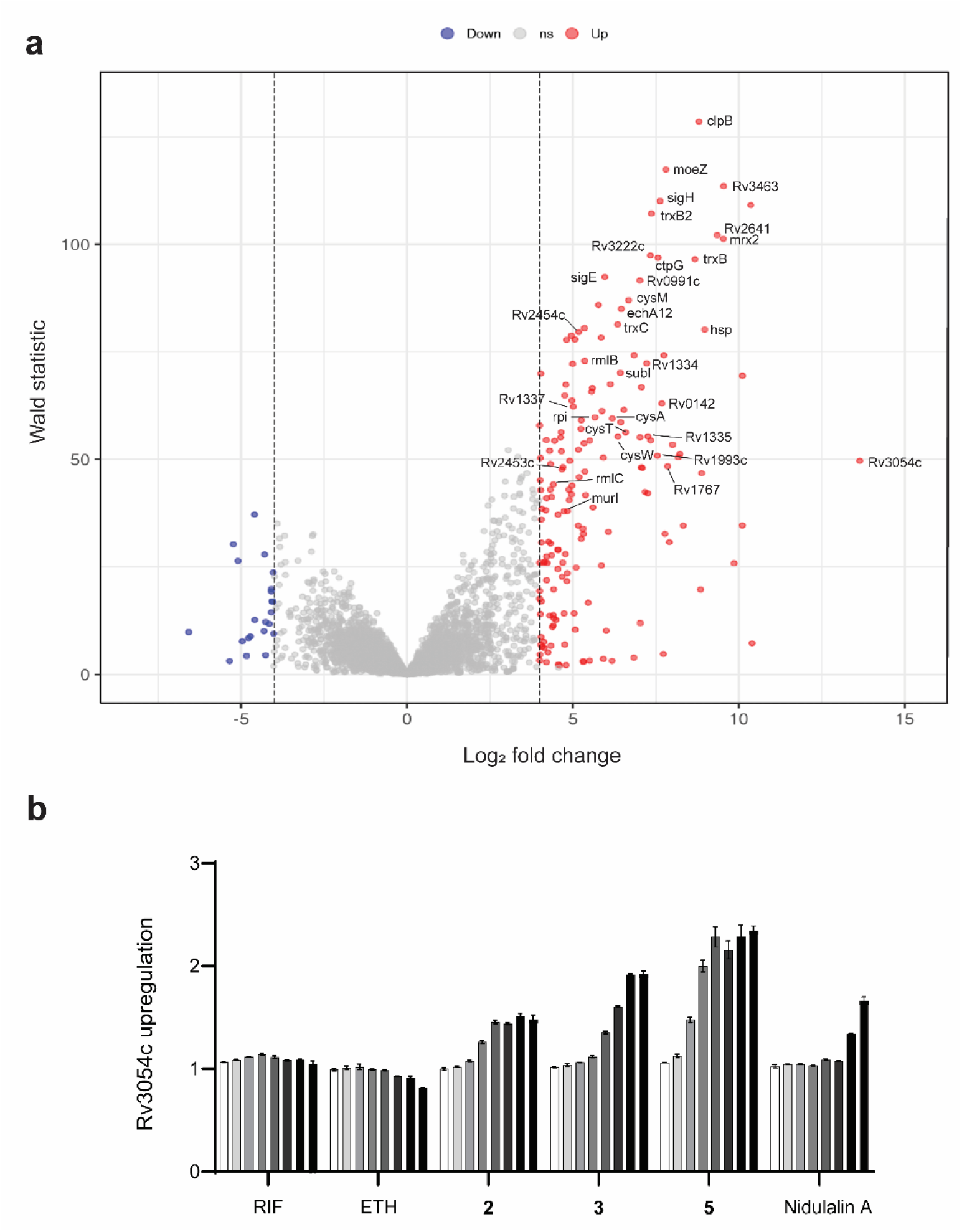
RNAseq data for *Mtb* treated with 10-fold the MIC of lead compound 2 for 2 hours. **a,** Members of the SigH regulon are labelled with gene names or Rv numbers for conserved hypotheticals of unknown function. **b,** Upregulation of Rv3054c expression using a promoter fusion previously described.

The strong upregulation of the SigH regulon was reminiscent of the transcriptional response to thiol-reactive mycotoxins we have previously reported^12^. We performed unsupervised clustering with our previously reported RNAseq datasets from the fungal mycotoxins nidulalin A and patulin (Fig. 6). We have previously shown that these mycotoxins kill *Mtb* cells by catastrophic thiol stress and key thiol stress response regulons are all clearly engaged by TTC mediated killing. Despite sharing a similar activation protein (EthA), *Mtb* treated with ethionamide and **2** showed no similarity in transcriptional responses.

**Fig. 6:**
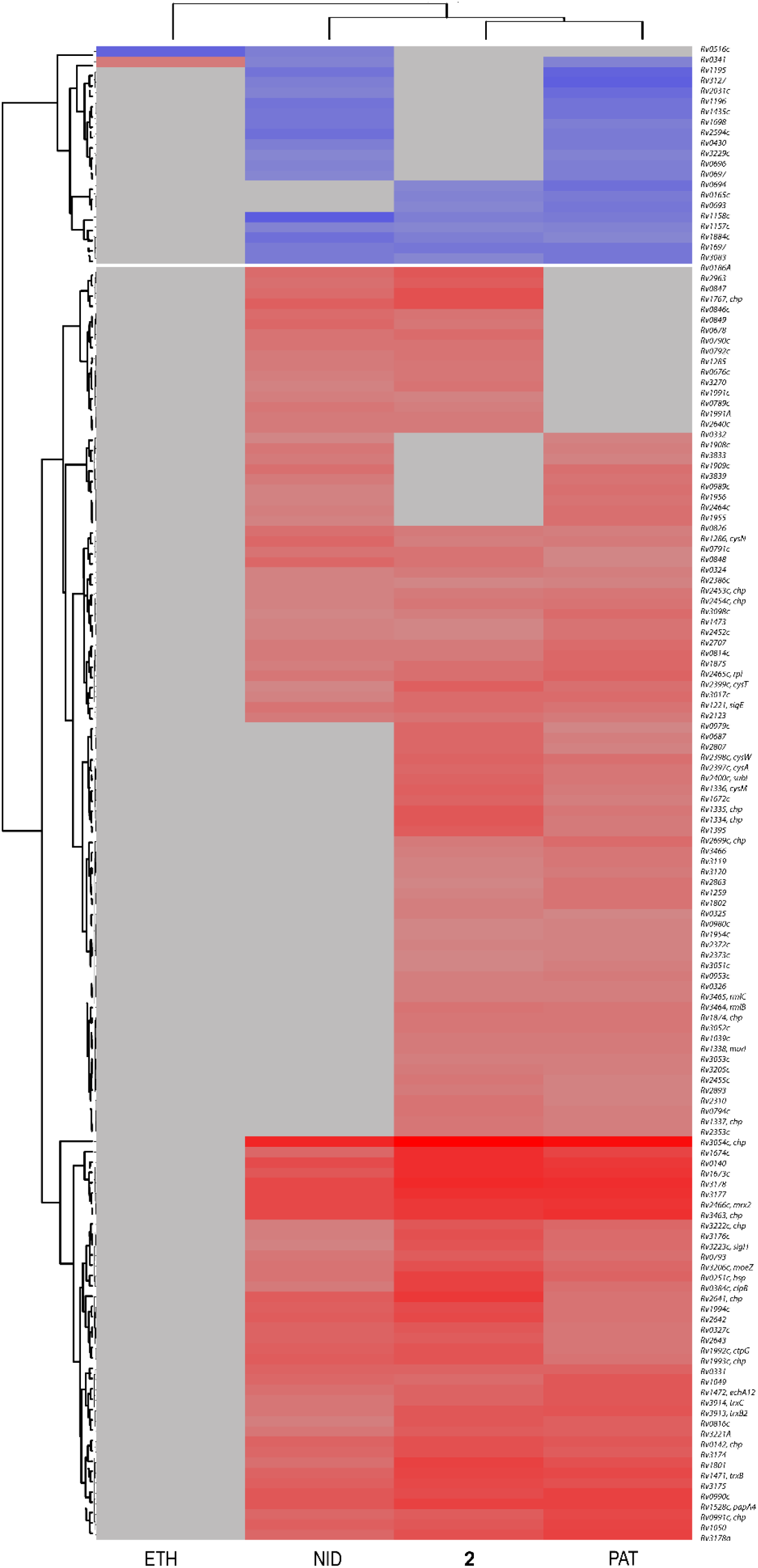
Clustering of transcriptional responses of *Mtb* to ethionamide (ETH), nidulalin A (NID), patulin (PAT) and lead compound 2. Hierarchical clustering of RNAseq data for the four indicated compounds showing **2** clusters closely with the fungal mycotoxins and shares no similarity with ETH. SigH family members are indicated by named genes in the list.

Rv3054c, a SigH regulon member that was the most strongly upregulated gene by all three treatments and has also previously been associated with thiol stress^36,37^. We previously constructed a reporter strain fusing the promoter region of this gene to mScarlet. Analysis of cells expressing this reporter confirmed upregulated expression of this locus upon treatment with **2** (Fig. 5b). Finally, we employed the reengineered redox sensitive GFP that monitors intracellular mycothiol status (Mrx1-roGFP2)^38^ and showed **2** treatment collapses mycothiol redox status (Supplementary Fig. 7).

## Discussion

Thiol homeostasis plays a critical role in the survival of *Mtb* to oxidative stress during infection and is primarily maintained by two low molecular weight thiols, mycothiol and ergothioneine. These two thiols play overlapping but distinct roles in evading damage from oxidative stress^39^. Mycothiol, like glutathione in mammalian systems, plays a unique role in detoxifying reactive electrophiles. The thiol in mycothiol can act as a nucleophile to trap a potential toxin and subsequent cleavage of that product by mycothiol conjugate amidase (encoded by *mca*) allows for excretion of the mercapturic acid captured product^40^. Chemical inhibition of mycothiol biosynthesis has been shown to be lethal for replicating and non-replicating cultures of *Mtb*^41,42^. Thiol redox status is also sensed by PknG which plays a role in survival in granulomas and under non-replicating conditions^43^. Thiol redox status also plays a role in determining expression of multiple cell envelope lipids involved in persistence through the sensor WhiB3^44^.

TTCs are rapidly cidal to *Mtb* cells. Initial oxidation of the thienothiazole core on the sulfur of the thiophene ring creates an initial electrophilic moiety that captures one equivalent of a cellular thiol. The product of this initial reaction rearranges to form an unusually stable sulfenic acid that then reacts with a second equivalent of a cellular thiol. Experiments with a ^14^C-labeled **4** failed to reveal a specific protein target with the majority of the label going into two active metabolites but there was also a diffuse labelling of many proteins consistent with wide reactivity with many cysteine residues across the proteome. Similarly, when we labelled cells with the photoaffinity probe there was broad labelling across the proteome without a specific protein target. Thus, inactivation of many proteins following reaction of exposed cysteines coupled with depletion of the intracellular protective thiol buffers mycothiol and ergothioneine results in a catastrophic thiol collapse. The transcriptional response is identical to the response of cells treated with the potent electrophilic mycotoxins nidulalin and patulin^12^. AC2P36 (5-chloro-N-(3-chloro-4-methoxyphenyl)-2-methylsulfonylpyrimidine-4-carboxamide) and AC2P20 (*N*-1,3-benzothiazol-2-yl-2-[(4,6-dioxo-5-phenyl-1,4,5,6-tetrahydropyrimidin-2-yl)thio]acetamide) have also been reported to kill *Mtb* by depletion of intracellular thiols and exposed *Mtb* cells show an extremely similar transcriptional profile to the TTCs^45^. Auranofin, a thiophilic organogold compound, has also been proposed to inhibit thiol redox homeostasis^46^.

A major goal of contemporary TB drug development is to create regimens that will markedly shorten the total duration of therapy. Phenotypically tolerant bacteria are thought to represent one of the major bottlenecks to shortening treatment times because such bacteria are non-replicating^47^. Most TB drugs have targets that are primarily sensitive to inhibition during active bacterial replication and thus contribute little to shortening^8^. Targeting processes such as thiol homeostasis that are important whether the bacteria is replicating or not presents the possibility for achieving real treatment shortening. The activity of the TTCs in non-replicating models of disease such as the starvation model in this study support that these agents may play an important role in regimens that are significantly shorter in duration.

The clinical use of ethionamide (or more commonly prothionamide) to treat multidrug- resistant (MDR) TB historically has created some level of existing resistance. Resistance to ethionamide occurs through either promoter mutations that upregulate the target *inhA* and less commonly through loss of the activator EthA. More contemporary regimens for MDR TB do not include this agent so the existing pool of strains that would have pre-existing resistance should be small. Alpibectir, an agent that induces expression of the monoxygenases activating ethionamide, has shown promising results in Phase 2 clinical trials^48,49^ and may overcome existing resistance to ethionamide. These “ethionamide boosters” may also be useful in combination with the molecules described in this study.

In many drug discovery programs for TB the transition from *in vitro* activity to *in vivo* activity is one of the most challenging aspects. This is likely attributable at least in part to targets that are often required for cell growth as opposed to targets that actively cause cell death. The delicate nature of cellular redox pools is highlighted by the fact that in nature fungal competitors of mycobacteria have independently evolved ways of targeting thiols as the most effective way to kill^12^. Unfortunately, fungal mycotoxins are impractical starting points for the development of useful TB drugs. There have been two compounds selected by screening for activity under acidic conditions that also cause depletion of cellular thiol pools^45,50^ but these have not been translated into agents suitable for *in vivo* studies. This series of TTC compounds was noteworthy for consistently performing better *in vivo* than *in vitro*, further emphasizing the importance of thiol redox balance during infection.

### Methods MIC

MIC measurements were performed as previously described^51^, in addition to the screening media, BSA free media was also used DPPC/Cas/Tx (4.7 g l^-1^ Middlebrook 7H9, 6 µM DPPC, 0.8 g l^-1^ NaCl, 0.03% casitone and 0.05% Tyloxapol) and 7H9/Glu/Cas/Tx (4.7 g l^-1^ Middlebrook 7H9 broth base, 4 g l^-1^ glucose, 0.8 g l^-1^ NaCl, 0.03% casitone and 0.05% Tyloxapol). The MIC were determined by visual inspection of the microtitre plates after 1- or 2-weeks growth at 37 °C with MIC recorded as the concentration that fully inhibited growth using an enlarging inverted mirror.

### Intramacrophage efficacy

Efficacy of the TTCs in killing intracellular *Mtb* growing in J774A.1 macrophages was performed as previously described^52^ with bacterial burden enumeration performed after 3 and 7 days of drug treatment.

### Bactericidal assays

The ability of the TTC compounds to kill actively replicating *Mtb* was performed as previously described^53^. The cidality of the compounds against starved non-replicating cells was performed as follows. *Mtb* H37Rv (25 ml) was grown to an OD_650nm_ of 0.4 in 7H9/GBSA (4.7 g l^-1^ 7H9 base, 5 g l^-1^ albumin fraction V, 4 g l^-1^ glucose, 0.81 g l^-1^ NaCl, and 0.05% tyloxapol). Cells were harvested by centrifugation and was washed three times in PBS. Cells were resuspended in 25 ml PBS/0.05% Tyloxapol in a 250 ml polystyrene roller bottle (Corning USA) and incubated at 37 °C in a rolling incubator (40 rpm). After 4 days, a single cell suspension was generated by filtration through a 5 μm methyl cellulose filter and cells resuspended to 10^7^ CFU ml^-1^ in PBS/0.05% Tyloxapol. Diluted cells were incubated in a 24w tissue culture plate (Corning USA) at 1 ml/well with vehicle control (DMSO), 0.1 μM rifampicin, 20 μM ethionamide or the TTC compounds at the desired concentrations. The plate was incubated at 37 °C in a sealed ziplock bag for 7 days after which appropriate dilutions in 7H9/GBSA were plated on Middlebrook 7H11/OADC agar media for CFU enumeration. The cidality of the TTC compounds against *Mtb* residing in caseum from infected rabbits was performed as previously described^17^.

### Reporter assays for cell wall damage, thiol stress and mycothiol redox responses

The assays were performed as previously described^12,19,38^.

### Raising resistant mutants

To raise resistant mutants against **3**, *Mtb* H37Rv cells (10^7^, 10^8^, and 10^9^) were plated on 7H11 ADGNTw (5 mg ml^-1^ BSA, 2 mg ml^-1^ glucose, 2 mg ml^-1^ glycerol, 0.85 mg ml^-1^ NaCl, 0.05% (v/v) Tween80) plates containing 1 μM or 5 μM **3** and incubated at 37 °C for 3 weeks. Individual colonies were then picked and grown in fresh 7H9 ADGNTw (no drug) prior to confirmation of resistance status by broth MIC testing. Genomic DNA of the mutants was isolated using a CTAB method and sequenced as previously described^54^. Sequencing reads were mapped to the *Mtb* H37Rv reference genome using BWA^55^. A custom script was used to extract genetic variants (SNPs and indels) by comparative analysis with the parental genome sequence, applying standard filters to exclude sites with low coverage (<10×) or heterogeneous base-calls (<70% purity). After the identification of the *ethA* gene as the main site for resistant mutations in initial strains, the genomic DNA from further resistant strains was isolated by Chelex extraction^56^. From this genomic DNA the *ethA* gene (rv3854c) was PCR amplified using locus-specific primers (Supplementary Table 1). PCR products were subsequently purified, subjected to Sanger sequencing (Macrogen USA), and obtained sequences compared to the reference H37Rv *ethA* sequence. In instances where no mutations in the *ethA* gene were detected, the *ethR* gene (rv3855) was similarly analysed (Supplementary Table 1).

### EthA overexpression strain construction

An over-expression plasmid for EthA was generated by cloning the full length rv3854c gene downstream of the *hsp60* promoter within a modified pMV361^57^ plasmid backbone (kanamycin resistance gene replaced with a nourseothricin^58^ resistance gene). The plasmid was subsequently introduced into *Mtb* H37Rv by electroporation^56^ and positive transformants selected following growth on nourseothricin-containing (25 µg ml^-1^) 7H11 medium for 3 weeks. Genomic DNA from individual clones was isolated as described earlier and the presence of the integrated plasmid confirmed by PCR, using plasmid-specific primers (Supplementary Table 1).

### EthA purification

*E. coli* BL21 DE3 cells pET28a(+)-His-EthA were grown in LB broth with kanamycin and riboflavin (50 µg ml^-1^ and 2 µM, respectively) to an OD ∼0.6 followed by overnight IPTG induction (1 mM) at 22 °C. Cells were harvested (3440×g for 10 min at 4 °C) and stored at −80°C. For purification of EthA, frozen cell pellets were thawed on ice for 30−45 minutes before addition of lysis buffer (50 mM Tris-HCl (pH 8.8), 100 mM KCl, 1 mM β-mercaptoethanol, 1% Triton100×, 50 µM FAD, Roche cOmplete^TM^ protease inhibitor cocktail, and lysozyme (25 mg l^-1^). Suspensions were incubated at 4 °C on a HulaMixer^TM^ sample mixer for 1 h followed by lysis via sonication for 15−25 min (Qsonica Q700 at 40% amplitude, 10 s ON, 30 s OFF) on ice. Lysed cells were centrifuged at 20,000×g for 20 min at 4 °C and the supernatant was collected. Imidazole was added to the supernatant to a final concentration of 10 mM before adding the Ni-NTA beads (2.5 ml beads/40 ml supernatant) equilibrated in wash buffer 1 (50 mM Tris-HCl, 100 mM KCl, 1 mM β-mercaptoethanol, 20 mM Imidazole, pH 8.8). The solution was incubated for 1 h at 4 °C on a HulaMixer to allow binding. After incubation the sample was loaded onto a gravity column (BioRad), and the beads were allowed to settle. After the initial lysate was run through, the column was washed with 6 column volumes of wash buffer 1 followed by 4 column volumes of wash buffer 2 (50 mM Tris-HCl, 100 mM KCl, 1 mM β-mercaptoethanol, 40 mM imidazole, pH 8.8). Finally, 1.5 column volumes of elution buffer (50 mM Tris-HCl, 100 mM KCl, 1 mM β-mercaptoethanol, 500 mM imidazole, 10% glycerol) were run through and fractions were collected. Fractions containing EthA were bright yellow and protein purity confirmed by SDS-PAGE. A 30 kDa MWCO concentrator was used to concentrate the protein and remove the imidazole from the final sample by centrifugation at 3000 rpm at 4 °C in storage buffer (50 mM Tris-HCl, 100 mM KCl, 10% glycerol) including 3−4 washes with storage buffer with a final EthA yield of 5 mg ml^-1^. The protein was aliquoted, flash frozen in liquid nitrogen and stored at −80°C for later use.

### Chemical proteomics with a bifunctional probe 6

Relevant *Mtb* strains (*ethA* frameshift mutant and/or *ethA* overexpression strain) were initially grown to an OD_600_ of 0.5 in 90 ml 7H9 GBSA (with 25 μg ml^-1^ Nourseothricin for *ethA* overexpression strain) before harvesting by centrifugation at 4000 rpm, washing once with PBST, and finally resuspending in 20 ml PBST. 5 ml aliquots of the final resuspension were transferred into 4 × 50 ml tubes. For the competition experiment (Supplementary Fig. 1c), active analog **7** (Supplementary Fig. 1a) was first added to each sample to either 0, 1.5, 6, or 30 μM final concentration. The final solvent (DMF) concentration was kept constant between all samples at 0.15% (v/v). After 6 h incubation at 37 °C, 10 μM of **6** was added and samples were left to incubate for a further 2 h. For Supplementary Fig. 1d, 5 μM of **6**, alone, was added to each sample and left to incubate at 37 °C for the specified time points. At the end of each incubation period, samples were harvested by centrifugation, washed twice with PBST, and finally resuspended in 1 ml PBS. Each sample was then split into 2 × 500 μl and transferred to individual wells of separate, flat bottomed, clear 24 well plates, on ice. One plate was left in the dark, while the other was exposed to UV light (Blak-Ray® B-100AP, 365 nm) for 10 min. Cells were then collected again by centrifugation and resuspended in 500 μl PBS with 1 mg ml^-1^ lysozyme and 1× protease inhibitor (Roche). Samples were transferred to 2 ml screw cap tubes containing ∼100 μl 0.1 mm acid-washed zirconia beads and frozen at −80°C prior to further processing. Once ready, samples were thawed, subjected to 4 × 45 s of bead beating at 6500 rpm (cooling on ice for 5 min between runs), and finally centrifuged at 13,000 rpm for 5 min at 4 °C. The soluble fraction was then passed through a 0.22 μm filter and protein concentration estimated using the BCA assay. Samples were then subjected to copper-catalyzed azide-alkyne cycloaddition reactions by combining, in 1.5 ml microfuge tubes, 40−50 μg protein extract, 1 mM CuSO_4_, 0.1 mM TBTA, 1 mM TCEP, and 40 μM Azide-fluor 545. Reactions were left in the dark at RT for 1 h prior to separation via SDS-PAGE. Labelled proteins were visualized using a Bio-Rad ChemiDoc Imaging System (Ex 532 nm, Em 580 nm), followed by Coomassie staining and visual analysis for assessing total protein loading.

### Carbon-14 isotope radiolabeled 4 incorporation assay

0.3 µM of **4** was added to 10 ml *Mtb* culture and incubated with shaking either for 2 h or 6 h. Culture were then centrifuged at 1000 rpm for 15 m. Methanol:chloroform:water (2:1:2, v/v/v) biphasic extraction was performed to separate the lipid, metabolites and proteins. 0.1 volume of zirconium beads were added to the solution and bead beating was performed for 5 cycles of 30 s with 1 min of cooling period in between each cycle. Subsequently an additional volume of chloroform was added to promote the separation of lipid containing organic phase from aqueous phase resulting in final concentration to 2:2:2 of methanol:chloroform:water. Lipid containing organic phase was washed with water and the aqueous phase containing metabolites and proteins were centrifuged at 13,000 rpm for 20 min. Finally, upper and lower metabolite phase was collected, and the interphase protein disc was dried in water bath and dissolved in water. Radioactivity associated with each fractions were then quantified using Beckman Coulter LS 6500 scintillation counter and counts per min were recorded.

### Metabolites extraction from *Mtb* cells and purification of 5-M62 and 5-M191

*Mtb* H37Rv was grown to OD_650nm_ of 1.0 in 7H9/Glu/Cas/Tx and 10 ml inoculated into 10 l fresh 7H9/Glu/Cas/Tx to OD_650nm_ of 0.4. Cells were harvested by centrifugation and washed in 800 ml 3% glycerol in PBS. The washed cells were resuspended in 1.85 l glucose/saline (4 g l^-1^ glucose/0.81 g l^-1^ NaCl) containing 75 μM of **5**. The cell suspension was split over two 3 l polycarbonate baffled flasks (Corning USA) incubated at 37 °C with gentle rotation (40 rpm). After 2 days, the entire suspension was poured into 1 l glass bottles containing 250 ml chloroform and vigorously shaken. After quenching the cells with chloroform, the mixture was filtered through a celite pad to remove cell debris, and the organic layer was separated. The aqueous layer was extracted with chloroform again. The combined organic layers were washed with brine, dried over MgSO_4_, and evaporated *in vacuo*. The major metabolites, **5-M62** and **5-M191** were purified by preparative TLC (Uniplate^TM^ silica gel, 20 × 20 cm, 1000 µm). For the separation of both metabolites, the eluent condition of ethyl acetate/hexane (3/2, v/v) was used, and then repurification of crude **5-M191** and crude **5-M62** were performed using ethyl acetate/hexane mixtures of 3:1 and 1:2 (v/v), respectively.

For labeling of cells with deuterated methionine, cells were grown up in 7H9/Glu/Cas/Tx as described above and 1.4 l of culture harvested and resuspended in 130 ml 7H9/Glu/Cas/Tx containing 1 mM isotopically labelled L-methionine (methyl-*d_3_*)(Cambridge Isotope Laboratories, Inc.). Cells were incubated in 500 ml polystyrene roller bottles (Corning USA) for 2 days after which cells were harvested by centrifugation, cells washed in 50 ml 10% glycerol in water and resuspended in 100 ml glucose/saline containing 0.3 mM L-methionine (methyl-*d_3_*) and 75 μM **5** in a 1 l glass bottle. The cell suspensions were incubated at 37 °C with gentle rotation (40 rpm) for 2 days followed by quenching with chloroform and metabolite extraction as described above.

### LC-MS analysis of metabolites

Samples were analyzed on an Agilent 1290 Infinity UPLC system coupled with an Agilent G6460C triple quadrupole mass selective detector (LC-QqQ). The UPLC was equipped with an Agilent EclipsePlus C18 2.1 mm × 50 mm 1.8 µm reverse phase column with flow rate of 0.8 mL/minute with a mobile phase consisting of two: water with 0.1% formic acid (A) and acetonitrile with 0.1% formic acid (B). A gradient was applied with initial composition of 5% acetonitrile to 95% over 8 min. The mass data were collected using an LC-QqQ in ESI+ mode, and spectra were monitored at 254 and 420 nm.

### Enzyme reaction with thiol quenching

Initial small-scale reactions consisted of 50 mM Tris-HCl (pH 7)/100 mM NaCl/200 µM NADPH/100 µM TTC analog/10 µM BSA/2.5 µM methylated *N*-acetyl cysteine with the enzyme reaction initiated by addition of 10 µM EthA enzyme in a final volume of 100 µl. Triplicate reactions were quenched with an equal volume of 50:50 methanol:acetonitrile solution at 0, 30, 60 and 120 min and precipitates removed by centrifugation prior to LC-MS analysis.

Large scale reactions consisted of a total volume of 23 ml in 50 mM Tris-HCl (pH 7)/ 100 mM NaCl/200 µM NADPH/300 µM **5**/10 µM enzyme/10 µM BSA/2.5 µM methylated *N*-acetyl cysteine with components in organic solvents (methylated *N*-acetyl cysteine in 70% ethanol and **5** in DMF) added dropwise with shaking. Reactions were run for 2 h followed by quenching with methanol/acetonitrile solution (1/1, v/v). The mixture centrifuged to remove precipitates and the supernatant analysed by TLC and LC-MS.

For the purification of intermediates **I1** and **I2** of the enzyme reaction with **5** (**5-I1** and **5-I2**), the supernatant was suspended in ethyl acetate and the organic layer collected after phase separation. The aqueous layer was re-extracted with ethyl acetate and the combined organic layers were washed with brine, dried over Na_2_SO_4_, and evaporated *in vacuo*. Each intermediate was purified by preparative TLC (Uniplate^TM^ silica gel, 20 × 20 cm, 1000 µm) with the eluent condition of ethyl acetate/hexane (9/1, v/v). I1 (R*f* = ∼0.3) exhibited a strong absorption on the TLC at 254 nm and **5-I2** consisted of two yellow spots located below **5-I1** (R*f* = 0.2−0.25). Chemical modification by dimedone quenching of **5-I1** and methylation of **5-I2** with iodomethane were monitored by LC-MS.

### *mshA* mutants generation

*mshA* mutants were generated using ORBIT (Oligonucleotide-mediated Recombineering followed by Bxb1 Integrase Targeting) previously reported^59^. Briefly, *Mtb* H37Rv strain expressing pKM427 and pKM461 was grown to OD ∼0.7 in the 7H9/OADC/Tween80 broth containing Zeocin (25 µg ml^-1^) and Kanamycin (20 µg ml^-1^). RecT expression was induced with Anhydrotetracycline (500 ng ml^-1^) for 8 h and the culture was further treated with 0.2 M Glycine overnight. Electrocompetent cells were prepared by washing with 10% Glycerol. Cells were electroporated with 1 µg of ORBIT oligonucleotide (Supplementary Table 1) consisting of the 48 bp *attP* site flanked on both sides with 70 bp of the *mshA* gene and 200 ng of pKM464, as the integrating plasmid, was electroporated (2.5 kV, 1000 Ω, 25 μF) into washed cells. The cells were recovered in 7H9/OADC/Tween80 broth for 72 h after which dilutions were then plated on 7H11 OADC agar plates with hygromycin (50 µg ml^-1^). The colonies were identified with PCR using flanking primers (Supplementary Table 1) (WT:1672 bp; Δ*mshA*: 3221 bp) and further verified with sequencing.

### *Mtb* RNA isolation for transcriptome analysis

*Mtb* (OD_650nm_ 0.2, 25 ml) were treated with ethionamide or TTC compounds at 0.1×, 1× and 10× the MIC concentration for 6 or 12 h with DMSO-treated cultures treated in parallel. At each time point, cells were rapidly harvested (3440×g, 5 min) and immediately resuspended in 1 mL TRIzol followed by bead beating using 0.1 mm silica/zirconia beads (BioSpec Products) (2 cycles at 7000 rpm, 45 sec in Roche MagnaLyzer). Chloroform (0.3 ml) was added for phase separation and the aqueous phase transferred to fresh tubes containing an equal volume of 70% ethanol. RNA was isolated using a RNeasy kit (Qiagen) following the manufacturer’s instructions including DNA removal with DNase I. Total RNA was used for cDNA library preparation and RNAseq as previously described^12^.

The NIH supercomputer Biowulf was used to compare raw RNA counts for each treatment group to DMSO controls. Two replicates per group were included in the analysis. Log2 Fold Change (log2FC), stats, and p-values were generated from the raw counts by running them through the DESeq2 (version 1.5.2) package in R. The following libraries were then downloaded into R for Gene Ontology, heatmap, upset plot, and clustering analysis of all treatment groups: readxl, dplyr, tidyr, pheatmap, UpSetR, RColorBrewer, clusterProfiler, enrichplot, and ggplot2. For clustered heatmap generation, a threshold of genes differentially expressed in ≥ 2 conditions with a log2FC ≥ 4 was used. To generate volcano plots, the package “BiocManager” was used to install “EnhancedVolcano”. Log2FC and ABS[10^(-stat)^] values were used to create volcano plots with log2FC as the X axis and Wald Statistic as the y-axis. Significance cutoffs shown are log2FC ≥ 4 and Wald Statistic ≥ 0.01078946722, corresponding to a p_value_ = 0.05.

