## Supplemental Information for "Thienothiazolocarboxamides: rapidly cidal agents causing catastrophic thiol collapse in *Mycobacterium tuberculosis*"

[n] *These authors contributed equally*

### [ Table of Contents ]

1. Supplementary Figures

Fig. 1 ........ S3

Fig. 2 ........ S4

Fig. 3 ........ S4

Fig. 4 ........ S5

Fig. 5 ........ S5

Fig. 6 ........ S6

Fig. 7 ........ S8

2. Supplementary Tables

Table 1 ........ S9

Table 2 ........ S10

3. X-ray crystallography ........ S10

4. Chemistry ........ S12

4-1. Characterization of **5-M191** ........ S12

4-2. Characterization of **5-M62** ........ S14

4-3. Synthesis of bifunctional chemical probe **6** ........ S17

4-4. Synthesis of radioactive $^{14}C$-labeled **4** ........ S20

4-5. Chemical oxidation test and confirmation of thiol adduct ........ S21

#### 1. Supplementary Figures

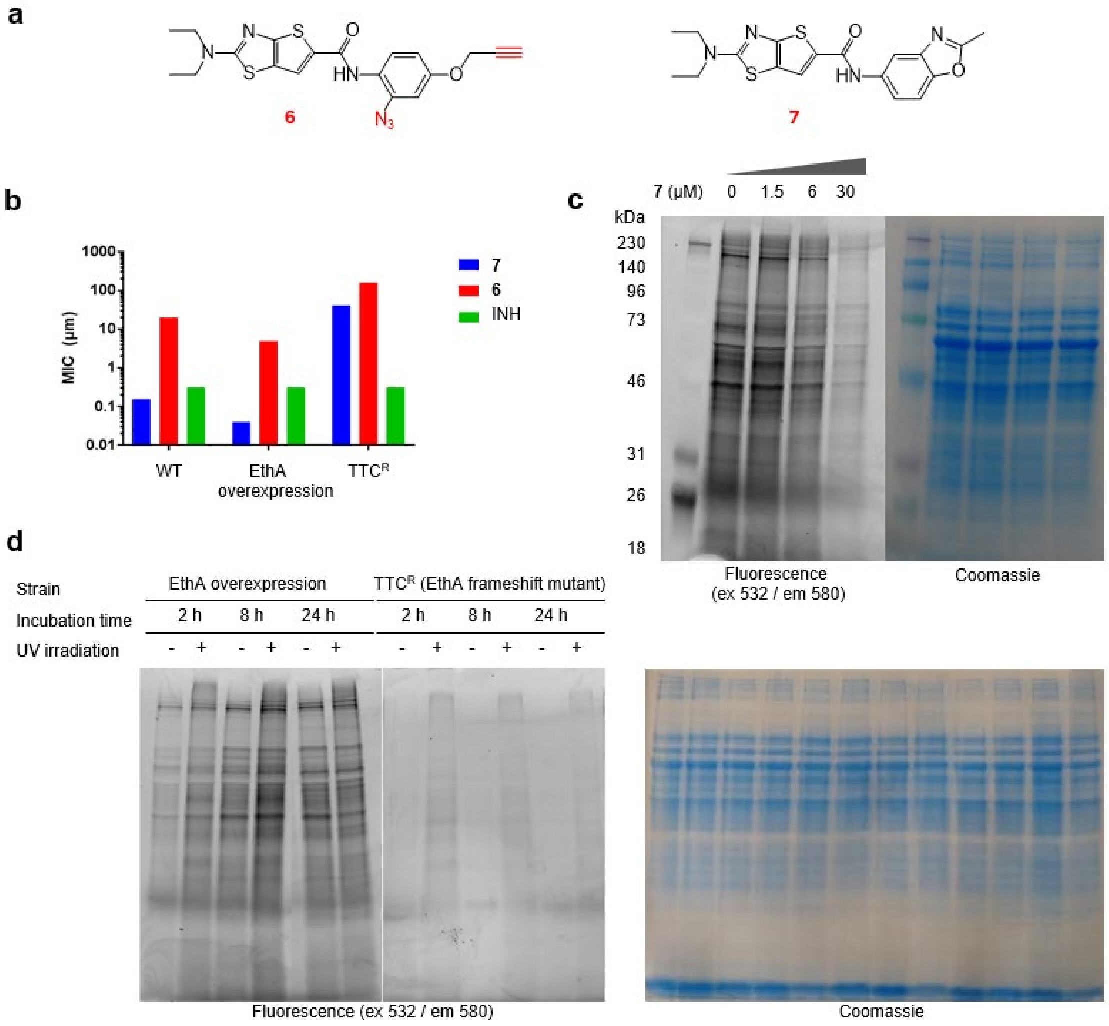

**Fig. 1. Chemical proteomics with a bifunctional probe 6 demonstrates lack of unique targets in *Mtb* despite EthA-mediated labeling. a,** The probe **6** contained a UV activatable azide group and a click chemistry linkable alkyne group and **7** is one of active TTC analogs. **b,** The click chemistry-**6** retained EthA-mediated activity against *Mtb* cells as evidenced by enhanced potency against an *ethA*-overexpressing strains contrasted by a 10-fold reduction in potency against an *ethA*-deficient strain. **c,** Protein labeling by 10 μM **6** can be outcompeted by **7**. **d,** Protein labeling is time-dependent but not UV-dependent although UV-activation increases protein labeling in treated cells. Protein labeling is dependent on EthA activity as evidenced by lack of labeling in an *ethA* frameshift mutant. Left panel shows fluorescence (Ex 532 nm/ Em 580 nm) in SDS-PAGE gels after Cu-catalyzed chemistry of cell lysate proteins with Azide-fluor 545. Right panel demonstrates protein profiles from lysates as revealed by Coomassie Blue staining.

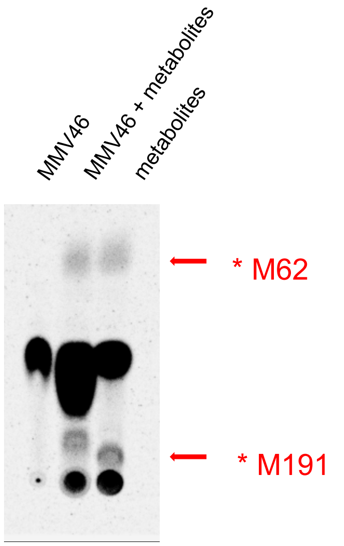

**Fig. 2. TLC analysis of compound 4-labeled free cytosolic metabolites in *Mtb*.** Cells were treated with $^{14}$C-labeled **4** and extracted metabolites separated by TLC followed by phosphor image analysis. The eluent condition was 1/1 of EA / Hexane.

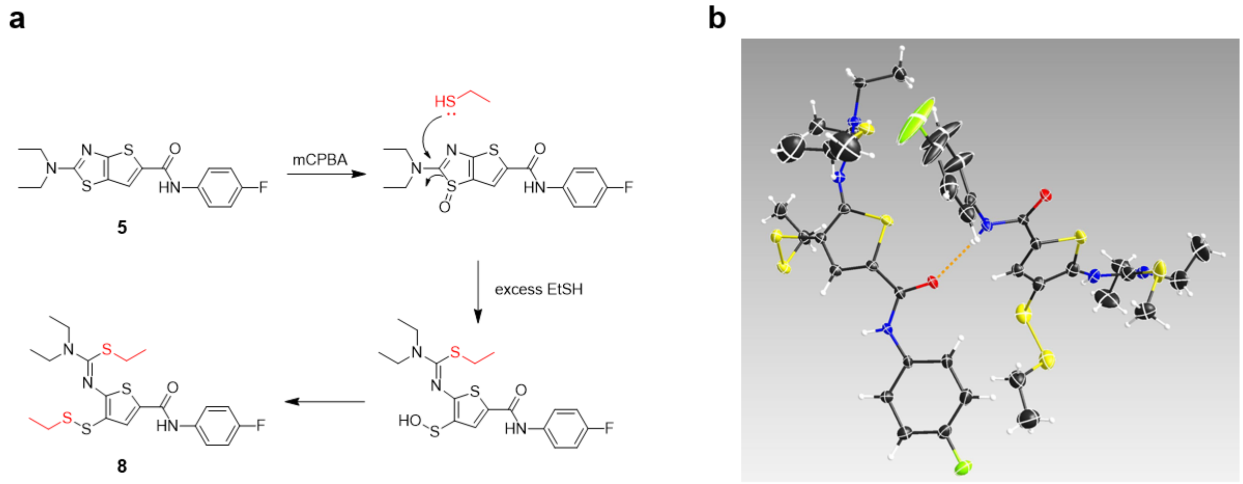

**Fig. 3. Chemical oxidation test of TTC analog**. **a,** Degradation scheme of **5** in the oxidation condition of mCPBA with EtSH quenching. **b,** Crystal structure of final adduct **8**.

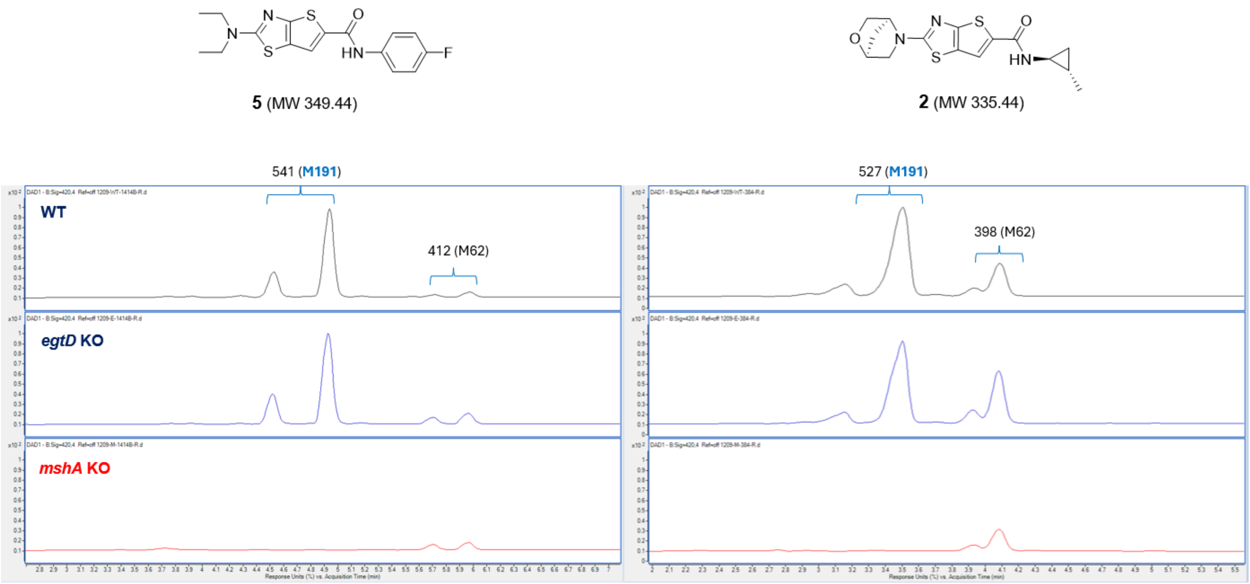

**Fig. 4. Chemical depletion of mycothiol by TTC analogs.** Major metabolites **M191** was not detected in *mshA* deletion strains, which is a glycosyltransferase catalyzing the first step in mycothiol biosynthesis. **M191** and **M62** were monitored by LC-MS at 420 nm in different strains and *egtD* deletion strains exhibited no effect.

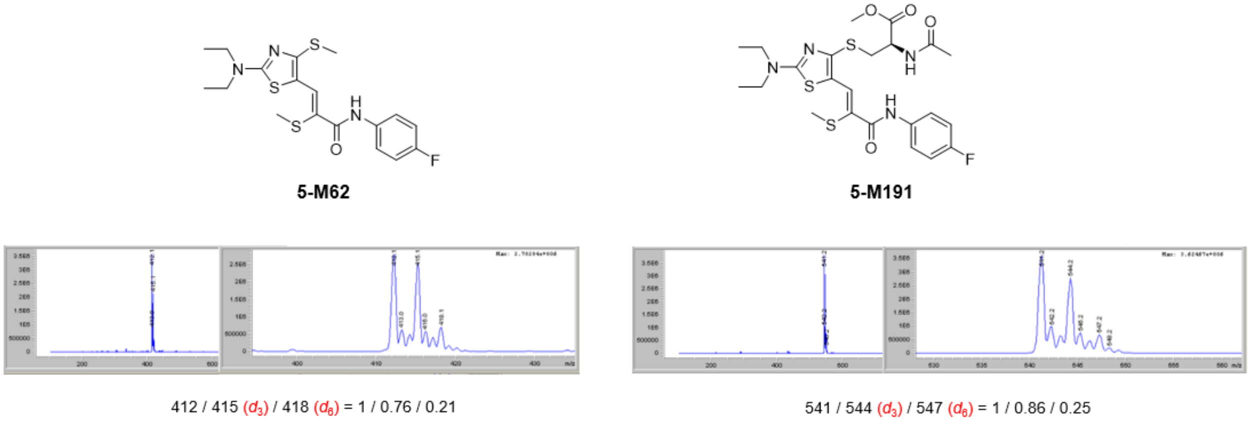

**Fig. 5. Confirmation of SAM-driven transmethylation in TTC metabolism.** The cells were incubated with **5** in the media containing stable isotope labeled methionine ($d_3$-Met) and metabolites, **M191** and **M62** were confirmed to embed labeled methyl groups by LC-MS.

**a**

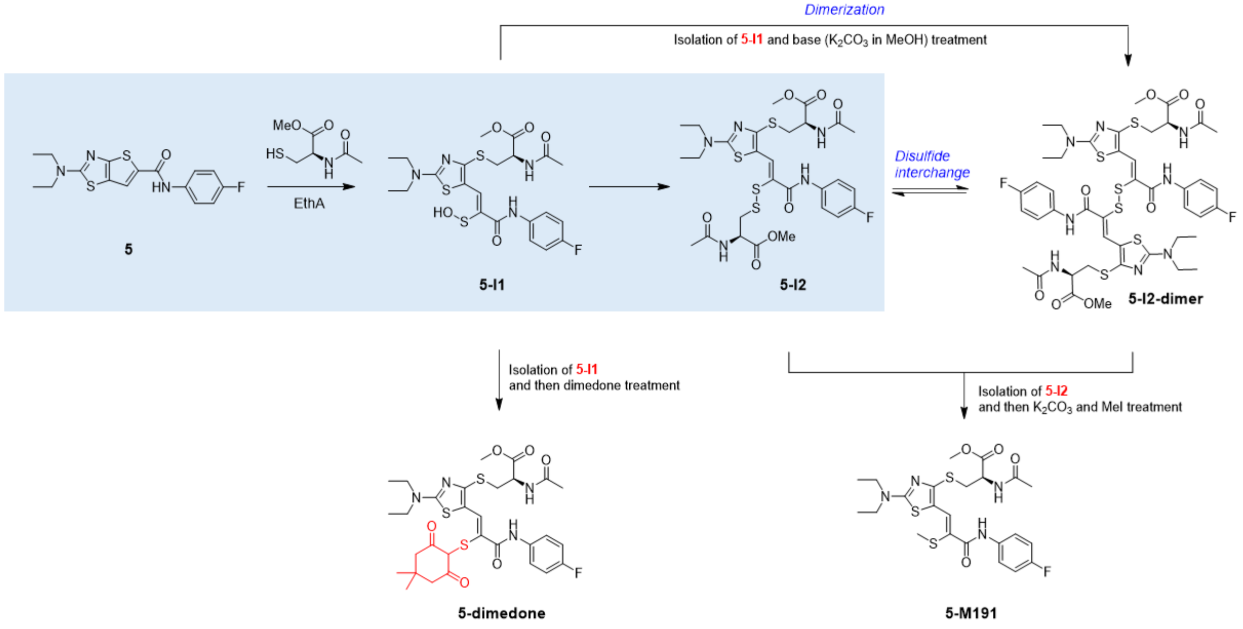
Dimerization
Isolation of 5-I1 and base ($K_2CO_3$ in MeOH) treatment
EthA
Disulfide interchange
5
5-I1
5-I2
5-I2-dimer
Isolation of 5-I1 and then dimedone treatment
Isolation of 5-I2 and then $K_2CO_3$ and MeI treatment
5-dimedone
5-M191

**b**

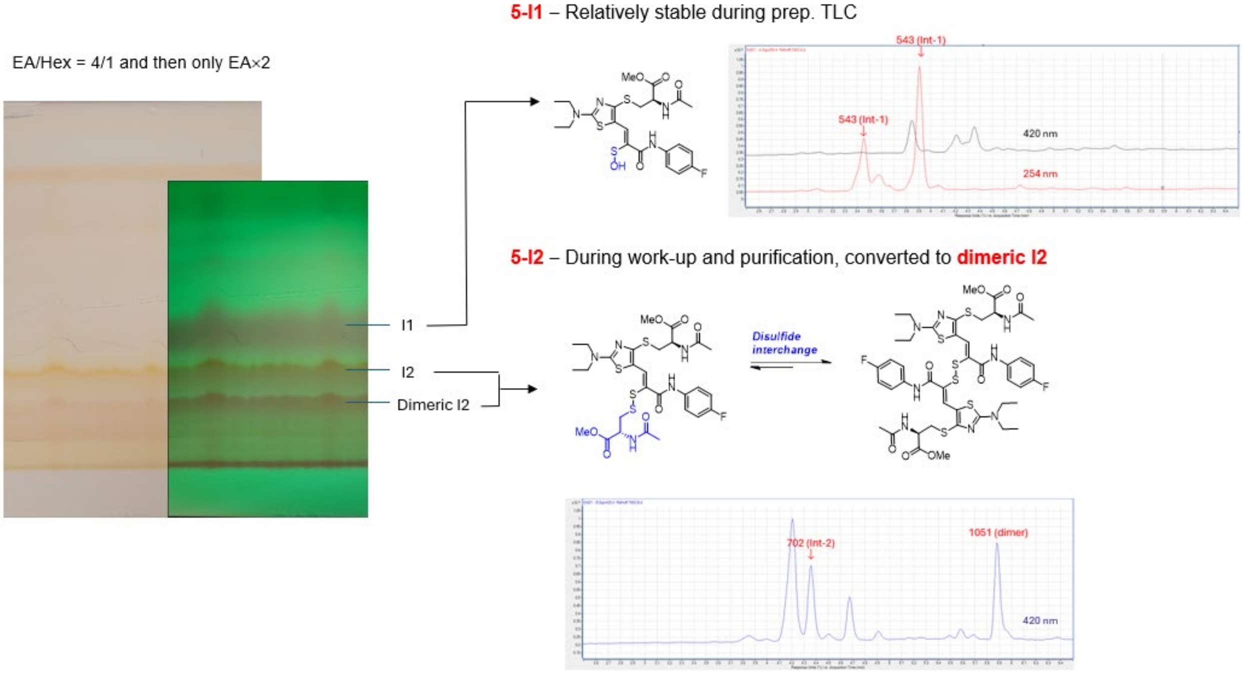
5-I1 – Relatively stable during prep. TLC
EA/Hex = 4/1 and then only EA×2
543 (Int-1)
543 (Int-1)
420 nm
254 nm
I1
I2
Dimeric I2
5-I2 – During work-up and purification, converted to dimeric I2
Disulfide interchange
702 (Int-2)
1051 (dimer)
420 nm

**c**

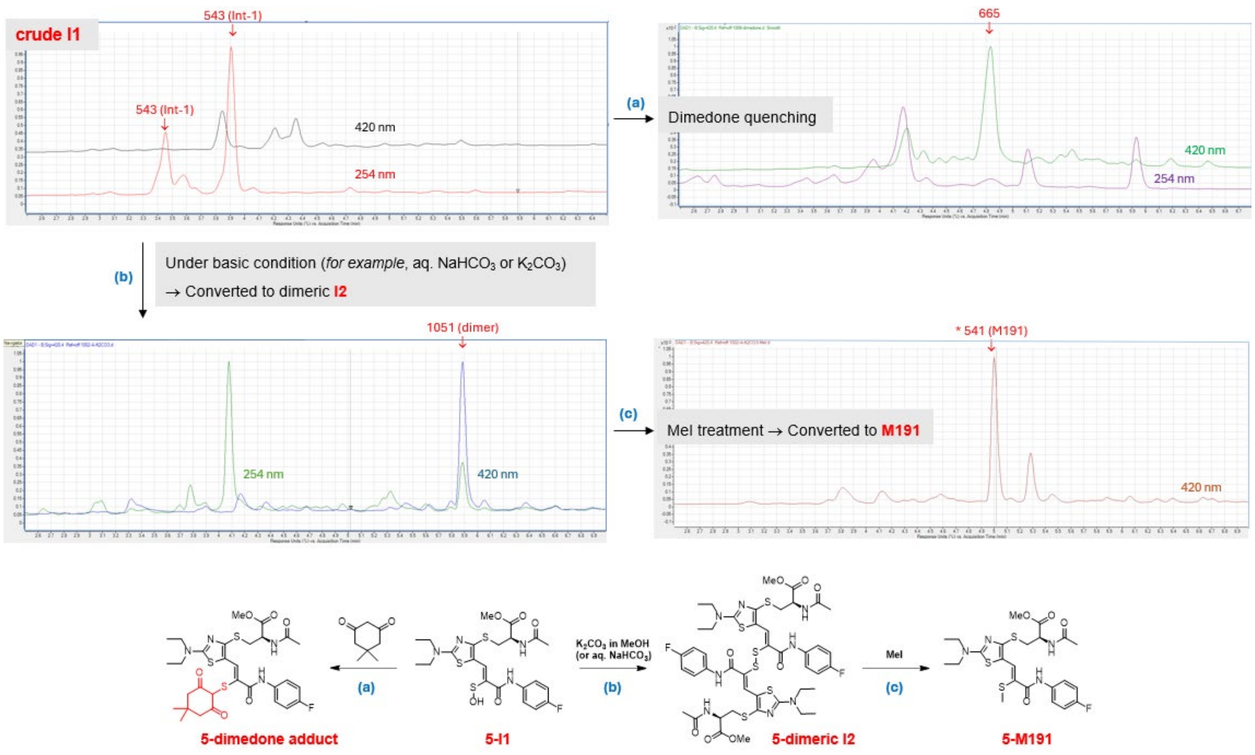

**d**

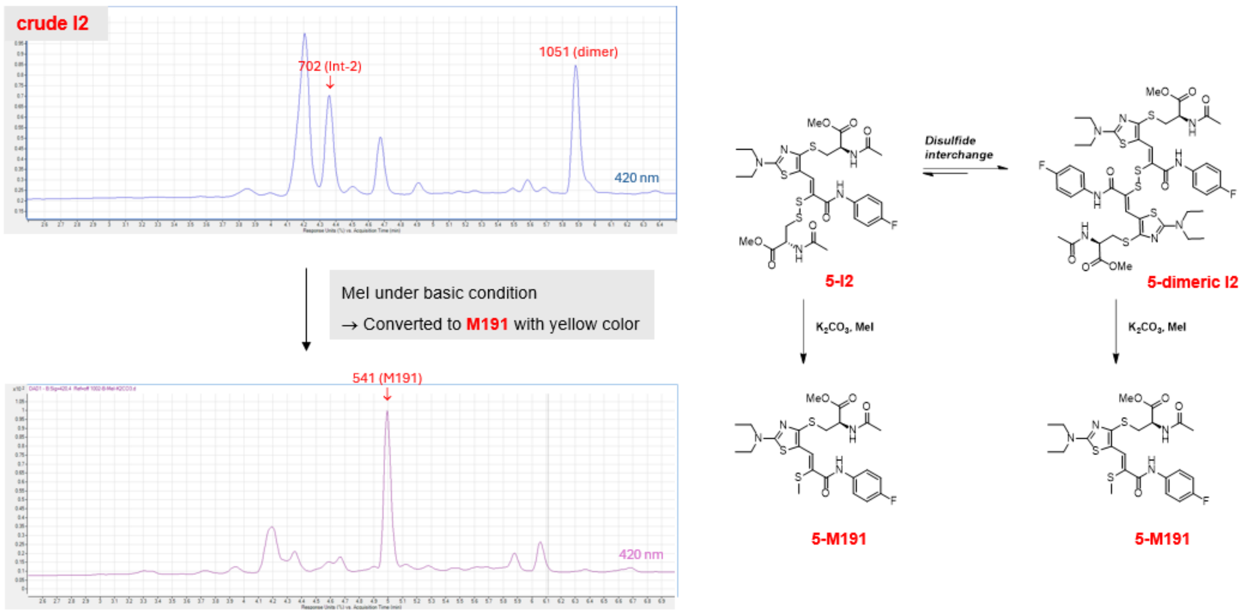

**Fig. 6. Enzymatic degradation in the presence of methylated *N*-acetyl cysteine. a,** Overall scheme of enzymatic and chemical degradation of **5**. **b,** Two major intermediates were purified by prep. TLC. **c, I1** was examined with chemical reactions to identify their structures. Sulfenic acid moiety of **I1** was confirmed by dimedone quenching. **d,** Disulfide **I2** was confirmed to transform the major metabolite **M191** when they were treated with iodomethane in the basic condition.

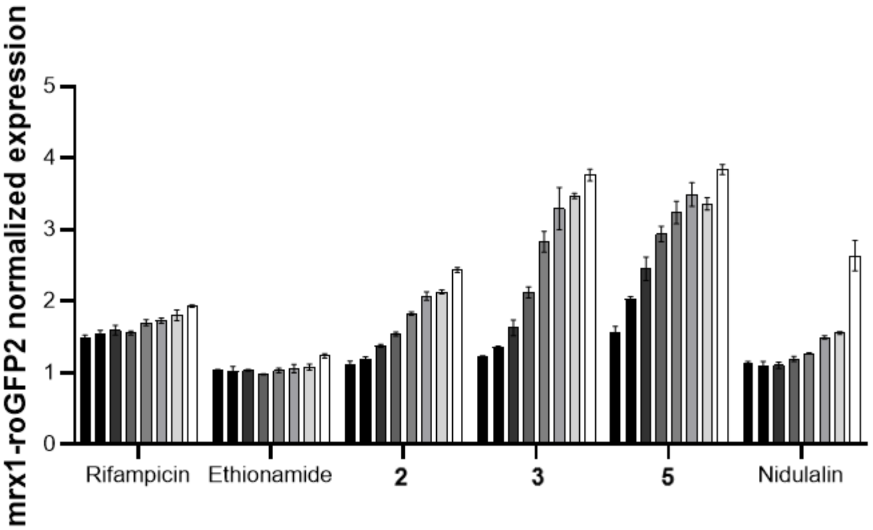

**Fig. 7. The TTC compounds oxidize the major redox buffer mycothiol in *Mtb*.** The mrx1-roGFP2 *Mtb* strain was exposed for 24 h to increasing concentrations of compounds (top concentrations of 5 μM for Rifampicin and 50 μM for ethionamide and compounds **2**, **3** and **5**) or nidulalin extract (0–50-fold MIC) and ratios of emission at 405/488 nm normalized to DMSO treated cells.

#### 2. Supplementary Table

**Table 1.** Oligonucleotides used in this study

| Oligonucleotide name | Sequence | Description |
|---|---|---|
| *ethA*-F | CACCCTGGCAGCTTACTACG | Forward primer for *ethA* sequencing |
| *ethA*-R | TTCGTTCCGGGCGATATCGC | Reverse primer for *ethA* sequencing |
| *ethR*-F | CGGAATCGGAAACCTAGCG | Forward primer for *ethR* sequencing |
| *ethR*-R | GGCTGCCAGATCGTCCAC | Reverse primer for *ethR* sequencing |
| *pMV361-ethA*-F | CAGCGAGGACAACTTGAGCC | Plasmid specific forward primer to verify *ethA* |
| *pMV361-ethA*-R | CGTTCGCCCTGTCGTTCACG | Plasmid specific reverse primer to verify *ethA* |
| *mshA*-ORBIT | CTGGCTGACCTCGAGCGCATTCTGGATCACACGTTGCACGGTCGGCAAGGAGGAAGTCACGCGCCCACCCGGTTTGTACCGTACACCACTGAGACCGCGGTGGTTGACCAGACAAACCATCGTGCCGCACACCTGCCATCCTTGCAGGAACCGAAGTGACACGCGCGGACGCGGCTCGTCAGCCGATC | Oligonucleotide for ORBIT knockout of *mshA* |
| *mshA*ins-F | GCAGGCATCGTGTCCCTAAGC | Primer 5’ to pKM464 insertion in *mshA* |
| *mshA*ins-R | TTGGGAGTACTTCAGCTGGCTG | Primer 3’ to pKM464 insertion in *mshA* |

#### 3. X-ray crystallography

All reflection intensities were measured at 110.00(10) K using a Rigaku XtaLAB Synergy R (equipped with a rotating-anode X-ray source and HyPix-6000HE detector) with Cu *K*α radiation (λ = 1.54178 Å) under the program CrysAlisPro (Version CrysAlisPro 1.171.42.49, Rigaku OD, 2022). The same program was used to refine the cell dimensions and for data reduction. The structure was solved with the program SHELXT-2018/2 (Sheldrick, 2018) and was refined on $F^2$ with SHELXL-2019/3 (Sheldrick, 2018). Analytical numeric absorption correction using a multifaceted crystal model was applied using CrysAlisPro. The temperature of the data collection was controlled using the system Cryostream 1000 from Oxford Cryosystems. The H atoms were placed at calculated positions (unless otherwise specified) using the instructions AFIX 23, AFIX 43 or AFIX 137 with isotropic displacement parameters having values 1.2 or 1.5 $U_{eq}$ of the attached C or N atoms.

**Table 2. Experimental details**

| | **5-M191** | **5-M62** |
|---|---|---|
| Crystal data | | |
| Crystal system, space group | Monoclinic, $P2_1$ | Triclinic, *P*-1 |
| Temperature (K) | 110 | 110 |
| *a*, *b*, *c* (Å) | 4.86749 (17), 17.0003 (8), 15.4848 (6) | 8.46914 (19), 8.7934 (2), 13.9587 (3) |
| $\alpha$, $\beta$, $\gamma$ (°) | -, 97.728 (4), - | 89.1348 (17), 73.7898 (19), 77.008 (2) |
| *V* (Å$^3$) | 1269.71 (9) | 971.40 (4) |
| *Z* | 2 | 2 |
| Radiation type | Cu *K*α | Cu *K*α |
| μ (mm$^{-1}$) | 3.06 | 3.68 |
| Crystal size (mm) | 0.08 × 0.04 × 0.03 | 0.24 × 0.15 × 0.12 |
| Data collection | | |
| Diffractometer | XtaLAB Synergy R, HyPix | |
| Absorption correction | Analytical<br>*CrysAlis PRO* 1.171.43.144a (Rigaku Oxford Diffraction, 2024)<br>Analytical numeric absorption correction using a multifaceted crystal model based on expressions derived by R.C. Clark & J.S. Reid. (Clark, R. C. & Reid, J. S. (1995). Acta Cryst. A51, 887-897)<br>Empirical absorption correction using spherical harmonics, implemented in SCALE3 ABSPACK scaling algorithm. | |
| $T_{min}$, $T_{max}$ | 0.838, 0.928 | 0.587, 0.766 |
| No. of measured, independent and observed [$I > 2\sigma(I)$] | 13843, 4137, 3644 | 31571, 3817, 3681 |

| | | |
|---|---|---|
| reflections | | |
| $R_{int}$ | 0.055 | 0.016 |
| (sin $\theta/\lambda)_{max}$ (Å$^{-1}$) | 0.598 | 0.616 |
| Refinement | | |
| $R[F^2 > 2\sigma(F^2)]$, $wR(F^2)$, $S$ | 0.042, 0.101, 1.03 | 0.028, 0.074, 1.05 |
| No. of reflections | 4137 | 3817 |
| No. of parameters | 327 | 463 |
| No. of restraints | 3 | 865 |
| H-atom treatment | H atoms treated by a mixture of independent and constrained refinement | H-atom parameters constrained |
| $\Delta\rho_{max}$, $\Delta\rho_{min}$ (e Å$^{-3}$) | 0.42, -0.40 | 0.29, -0.19 |
| Absolute structure | Flack x determined using 1397 quotients [(I+)-(I-)]/[(I+)+(I-)] (Parsons, Flack and Wagner, Acta Cryst. B69 (2013) 249-259). | - |
| Absolute structure parameter | 0.005 (16) | - |

Computer programs: *CrysAlis PRO* 1.171.42.49 (Rigaku OD, 2022), *SHELXT2018*/2 (Sheldrick, 2018), *SHELXL2019*/3 (Sheldrick, 2018), *SHELXTL* v6.10 (Sheldrick, 2008).

#### 4. Chemistry

$^1$H and $^{13}$C NMR spectra were recorded on a Varian Mercury-300 NMR Spectrometer and 2D NMR spectra on a Varian Mercury-600 NMR Spectrometer. The chemical shifts were measured in ppm relative to chloroform solvent signal (δ 7.26) or DMSO (δ 2.50). Routine mass analyses (LRMS) were performed on HP Agilent LC/MS series 1100 system equipped with a reverse phase column (Agilent Poroshell 120 EC-C18, 2.7 μm, 50 × 2.1 mm) and photodiode array detector using electrospray ionization (ESI). Accurate masses (HRMS) were obtained using Waters LCT Premiere time-of-flight mass spectrometer. The instrument was operated in the W-mode at a resolution of 10,000 and in positive ion mode. Ions were generated with the Z-Spray electrospray ionization (ESI) with a capillary voltage of 3.4 kV. Accurate masses were determined using the internal standard method. The metabolite was monitored using thin-layer chromatography (TLC) (silica gel 60 F254 0.25 mm), and the products were visualized by UV light (254 nm). SiliaFlash® P60 (40–60 μm) used in flash column chromatography was purchased from Silicycle Inc.

##### 4-1. Characterization of 5-M191

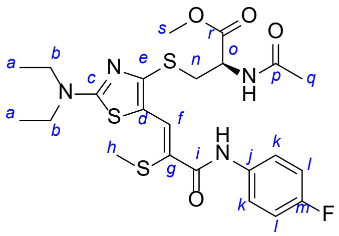

Yellow solid: $^1$H NMR (300 MHz, DMSO-$d_6$) δ 9.83 (s, 1H, *i*-NH-*j*), 8.47 (d, *J* = 7.7 Hz, 1H, *o*-NH-*p*), 8.05 (s, 1H, *f*-H), 7.77–7.72 (m, 2H, 2*k*-H), 7.19–7.13 (m, 2H, 2*l*-H), 4.61–4.54 (m, 1H, *o*-H), 3.74–3.68 (m, 1H, *n*-H), 3.61 (s, 3H, *s*-H3), 3.59–3.54 (m, 4H, 2*b*-H2), 3.34–3.24 (m, 1H, *n*-H), 2.23 (s, 3H, *h*-H3), 1.83 (s, 3H, *q*-H3), 1.24–1.19 (m, 6H, 2*a*-H3); $^{13}$C NMR (75 MHz, DMSO-$d_6$) δ 171.1 (*p*), 170.3 (*c*), 169.5 (*r*), 163.9 (*i*), 159.9 (*m*), 156.7 (*m*), 155.2 (*e*), 135.4 (d, *j*), 133.6 (*g*), 122.4 (d, *k*), 116.9 (*d*), 115.1 (d, *l*), 113.0 (*f*), 52.4 (*o*), 52.1 (*s*), 45.3 (*b*), 33.2 (*n*), 22.3 (*q*), 19.3 (*h*), 12.4 (*a*); HRMS (ESI) m/z calc'd for $C_{23}H_{30}FN_4O_4S_3$ [M + H]$^+$ 541.1413, found 541.1412.

*[NMR Spectra of **5-M191**]*

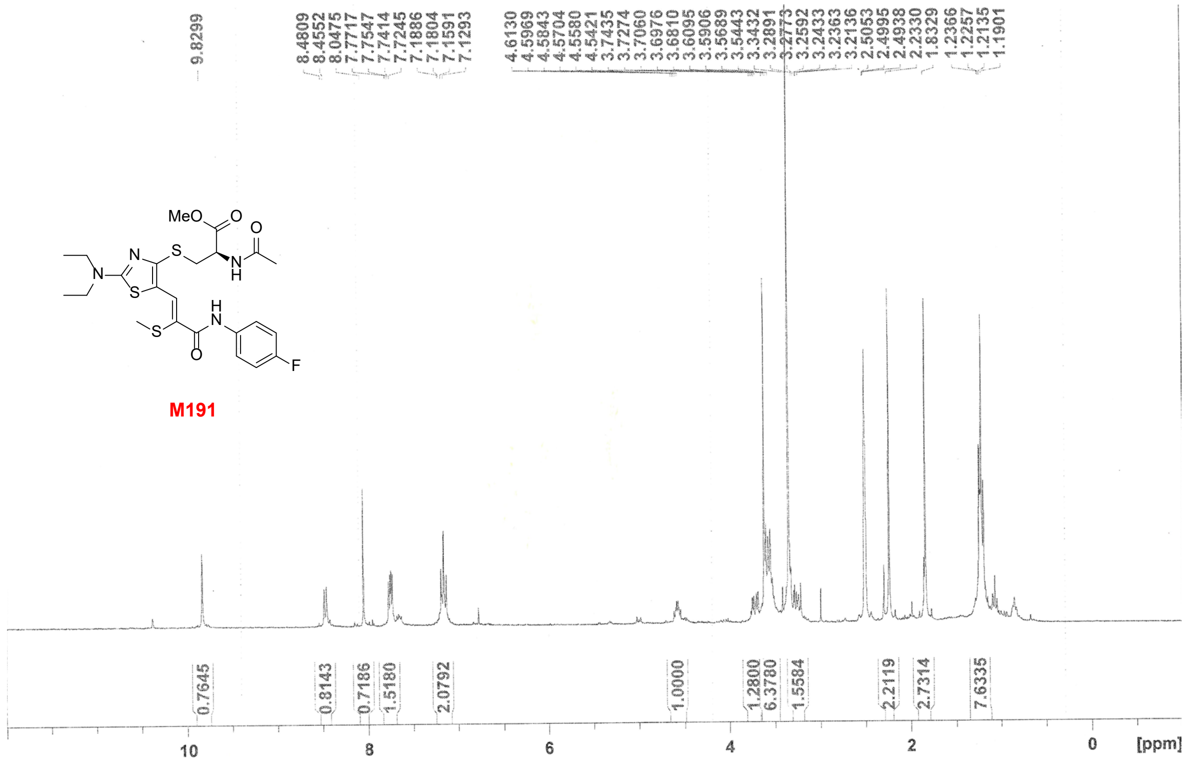
MeO
O
O
S
N
N
H
N
S
S
H
N
O
F
M191
9.8299
8.4809
8.4552
8.0475
7.7717
7.7547
7.7414
7.7245
7.1886
7.1804
7.1591
7.1293
4.6130
4.5969
4.5843
4.5704
4.5580
4.5421
3.7435
3.7274
3.7060
3.6976
3.6810
3.6095
3.5906
3.5689
3.5443
3.3432
3.2891
3.2773
3.2592
3.2433
3.2363
3.2136
2.5053
2.4995
2.4938
2.2330
1.8329
1.2366
1.2257
1.2135
1.1901
0.7645
0.8143
0.7186
1.5180
2.0792
1.0000
1.2800
6.3780
1.5584
2.2119
2.7314
7.6335
10
8
6
4
2
0
[ppm]

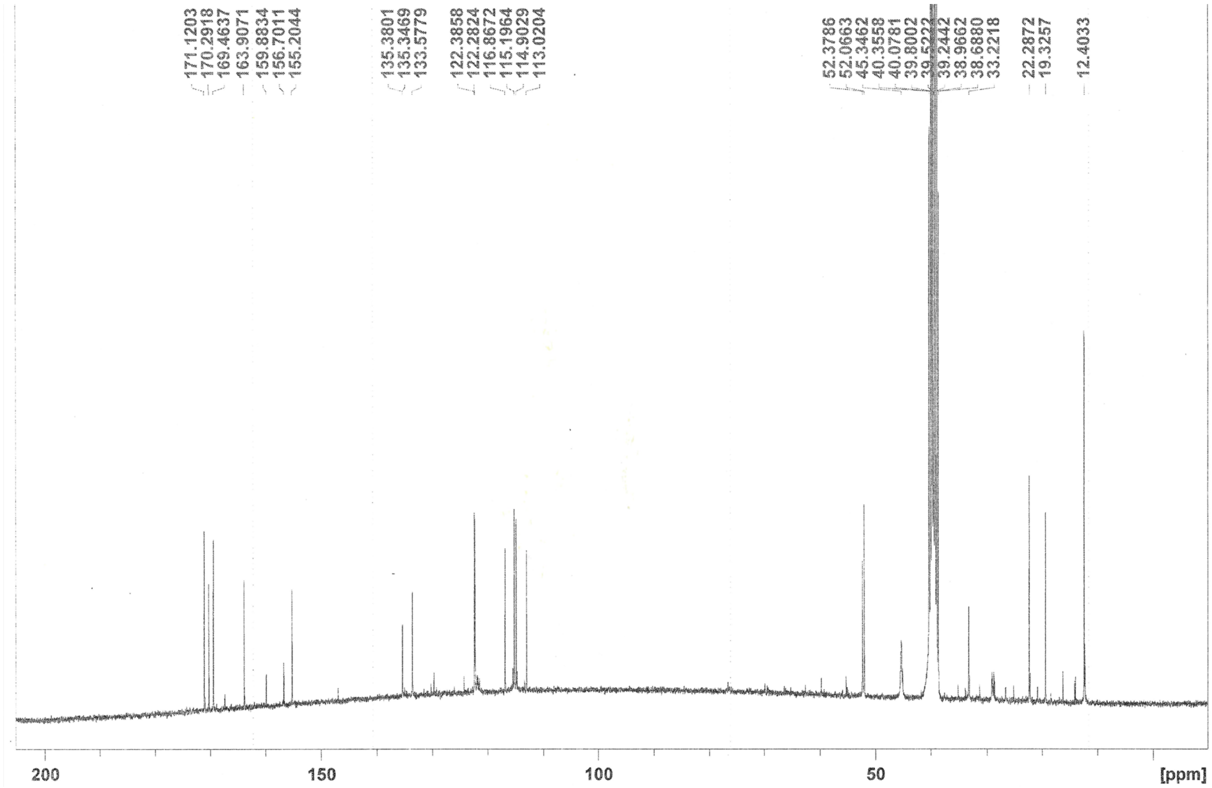
171.1203
170.2918
169.4637
163.9071
159.8834
156.7011
155.2044
135.3801
135.3469
133.5779
122.3858
122.2824
116.8672
115.1964
114.9029
113.0204
52.3786
52.0663
45.3462
40.3558
40.0781
39.8002
39.5222
39.2442
38.9662
38.6880
33.2218
22.2872
19.3257
12.4033
200
150
100
50
[ppm]

*[HRMS Spectra of **5-M191**]*

**Elemental Composition Report** **Page 1**

**Single Mass Analysis**
Tolerance = 5.0 mDa / DBE: min = -1.5, max = 100.0
Element prediction: Off
Number of isotope peaks used for i-FIT = 3

Monoisotopic Mass, Even Electron Ions
158 formula(e) evaluated with 3 results within limits (up to 50 closest results for each mass)
Elements Used:
C: 0-200 H: 0-200 N: 4-4 O: 0-20 32S: 2-3 F: 1-1

SO-10JAN25-MMV46-M191 155 (2.639) AM2 (Ar,22000.0,0.00,0.00); ABS
TOF MS ES+

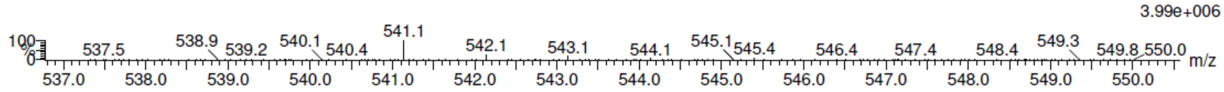

| | | | | |
|---|---|---|---|---|
| Minimum: | | | | -1.5 |
| Maximum: | | 5.0 | 5.0 | 100.0 |

| Mass | Calc. Mass | mDa | PPM | DBE | i-FIT | Norm | Conf(%) | Formula |
|---|---|---|---|---|---|---|---|---|
| 541.1412 | 541.1413 | -0.1 | -0.2 | 10.5 | 414.8 | 1.196 | 30.23 | C23 H30 N4 O4 32S3 F |
| | 541.1438 | -2.6 | -4.8 | 6.5 | 414.7 | 1.084 | 33.83 | C19 H30 N4 O9 32S2 F |
| | 541.1380 | 3.2 | 5.9 | 15.5 | 414.6 | 1.023 | 35.94 | C26 H26 N4 O4 32S2 F |

#### 4-2. Characterization of 5-M62

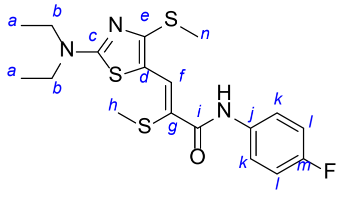

Yellow solid: [1]H NMR (300 MHz, DMSO-$d_6$) δ 9.81 (s, 1H, *i*-NH-*j*), 8.12 (s, 1H, *f*-H), 7.77–7.73 (m, 2H, 2*k*-H), 7.19–7.13 (m, 2H, 2*k*-H), 3.55 (q, *J* = 7.0 Hz, 4H, 2*b*-H2), 2.60 (s, 3H, *n*-H3), 2.23 (s, 3H, *h*-H3), 1.22 (t, *J* = 7.0 Hz, 6H, 2*a*-H3); [13]C NMR (75 MHz, DMSO-$d_6$) δ 170.3 (*c*), 164.0 (*i*), 159.9 (*m*), 157.5 (*e*), 156.7 (*m*), 135.4 (d, *j*), 133.9 (*g*), 122.3 (d, *k*), 116.3 (*d*), 115.0 (d, *l*), 112.4 (*f*), 45.3 (*b*), 19.4 (*h*), 15.2 (*n*), 12.4 (*a*); HRMS (ESI) m/z calc'd for $C_{18}H_{23}FN_3OS_3$ [M + H]$^+$ 412.0987, found 412.0984.

*[NMR Spectra of **5-M62**]*

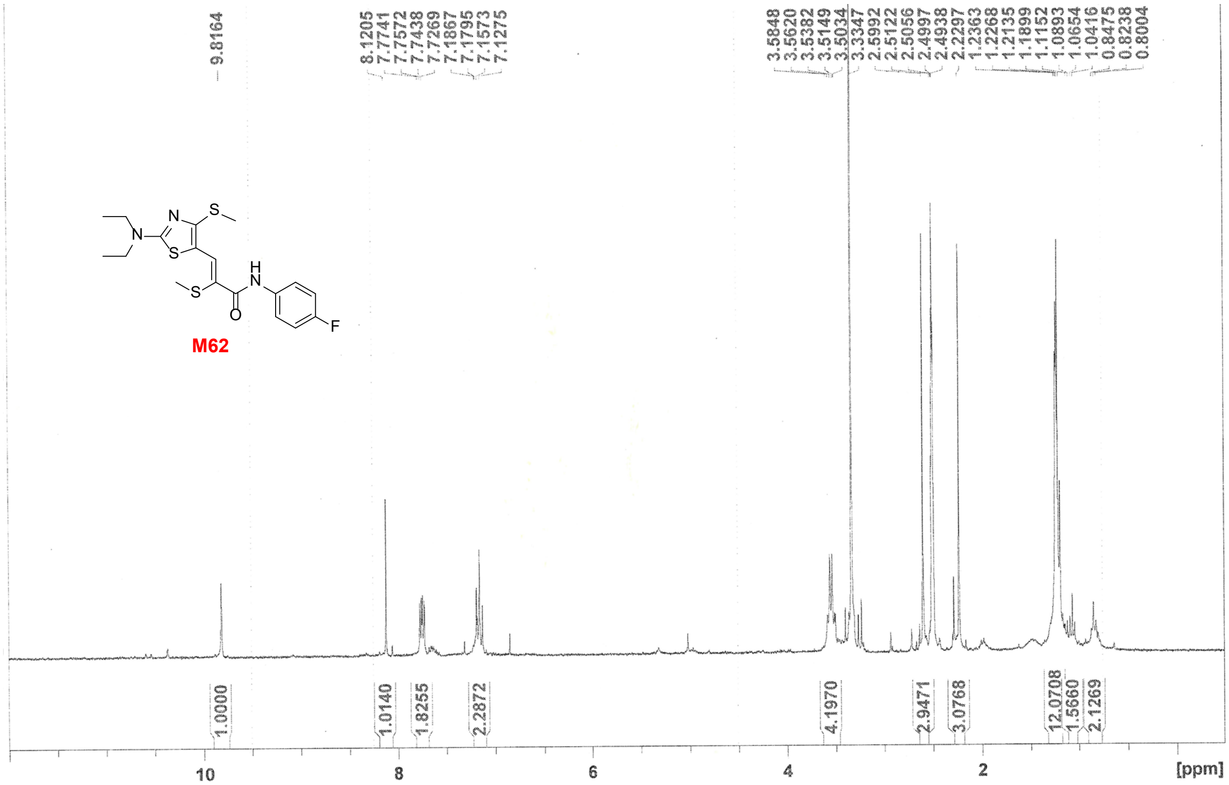

**M62**

9.8164
8.1205
7.7741
7.7572
7.7438
7.7269
7.1867
7.1795
7.1573
7.1275
3.5848
3.5620
3.5382
3.5149
3.5034
3.3347
2.5992
2.5122
2.5056
2.4997
2.4938
2.2297
2.2363
1.2268
1.2135
1.1899
1.1152
1.0893
1.0654
1.0416
0.8475
0.8238
0.8004
1.0000
1.0140
1.8255
2.2872
4.1970
2.9471
3.0768
12.0708
1.5660
2.1269
10
8
6
4
2
[ppm]

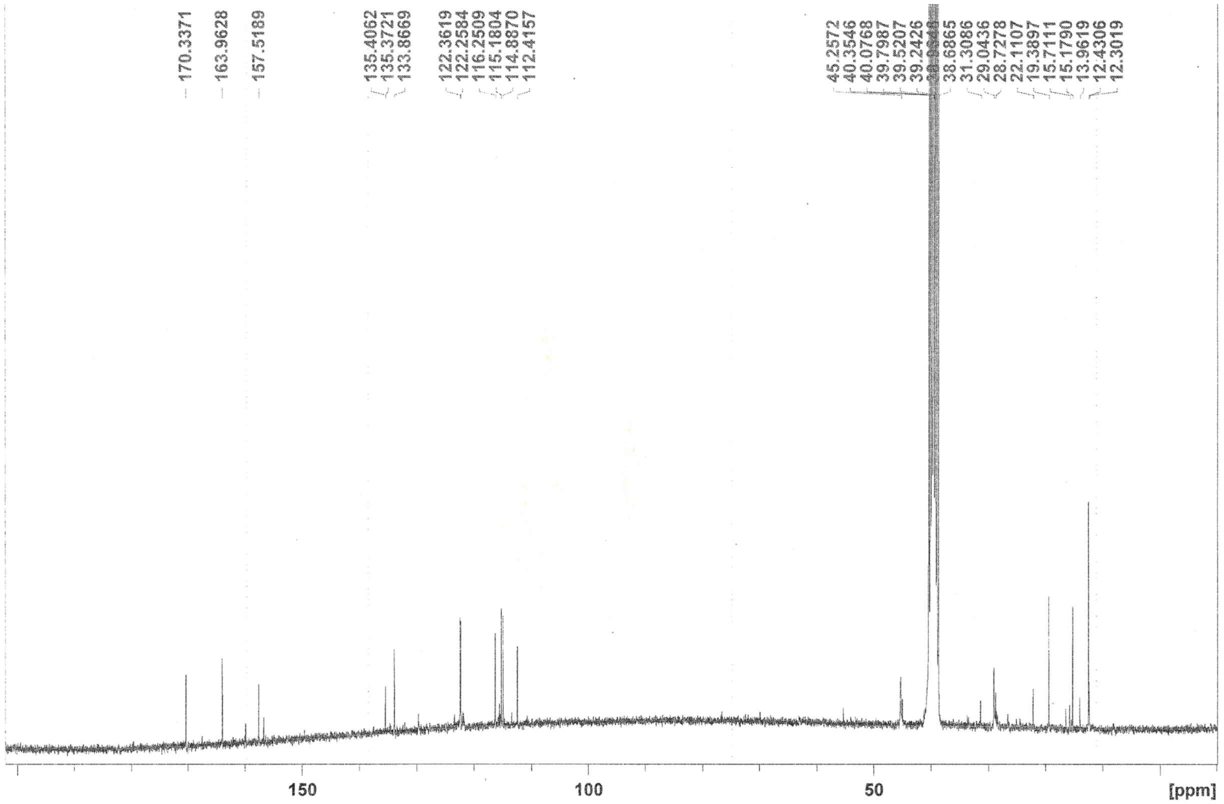
170.3371
163.9628
157.5189
135.4062
135.3721
133.8669
122.3619
122.2584
116.2509
115.1804
114.8870
112.4157
45.2572
40.3546
40.0768
39.7987
39.5207
39.2426
38.6865
31.3086
29.0436
28.7278
22.1107
19.3897
15.7111
15.1790
13.9619
12.4306
12.3019
150
100
50
[ppm]

*[HRMS of **5-M62**]*

#### Elemental Composition Report

##### Single Mass Analysis

Tolerance = 3.0 PPM / DBE: min = -1.5, max = 100.0
Element prediction: Off
Number of isotope peaks used for i-FIT = 3

Monoisotopic Mass, Even Electron Ions
3781 formula(e) evaluated with 4 results within limits (up to 50 closest results for each mass)
Elements Used:
C: 15-19 H: 0-200 N: 0-10 O: 0-40 F: 0-2 32S: 0-4
SO-27FEB25-MMV46-M62 110 (1.877) AM2 (Ar,22000.0,0.00,0.00); ABS
TOF MS ES+

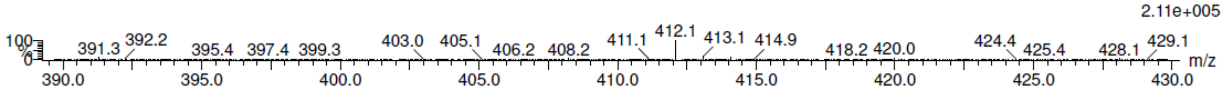

| | | | | | | | | |
|---|---|---|---|---|---|---|---|---|
| Minimum: | | | | -1.5 | | | | |
| Maximum: | | 5.0 | 3.0 | 100.0 | | | | |
| Mass | Calc. Mass | mDa | PPM | DBE | i-FIT | Norm | Conf(%) | Formula |
| 412.0984 | 412.0987 | -0.3 | -0.7 | 8.5 | 451.6 | 2.199 | 11.10 | C18 H23 N3 O F 32S3 |
| | 412.0979 | 0.5 | 1.2 | 9.5 | 450.6 | 1.173 | 30.95 | C17 H19 N3 O6 F 32S |
| | 412.0992 | -0.8 | -1.9 | 9.5 | 450.3 | 0.884 | 41.30 | C16 H18 N3 O10 |
| | 412.0992 | -0.8 | -1.9 | 14.5 | 451.2 | 1.793 | 16.64 | C18 H15 N7 O2 F 32S |

#### 4-3. Synthesis of bifunctional chemical probe 6

[Synthetic scheme]

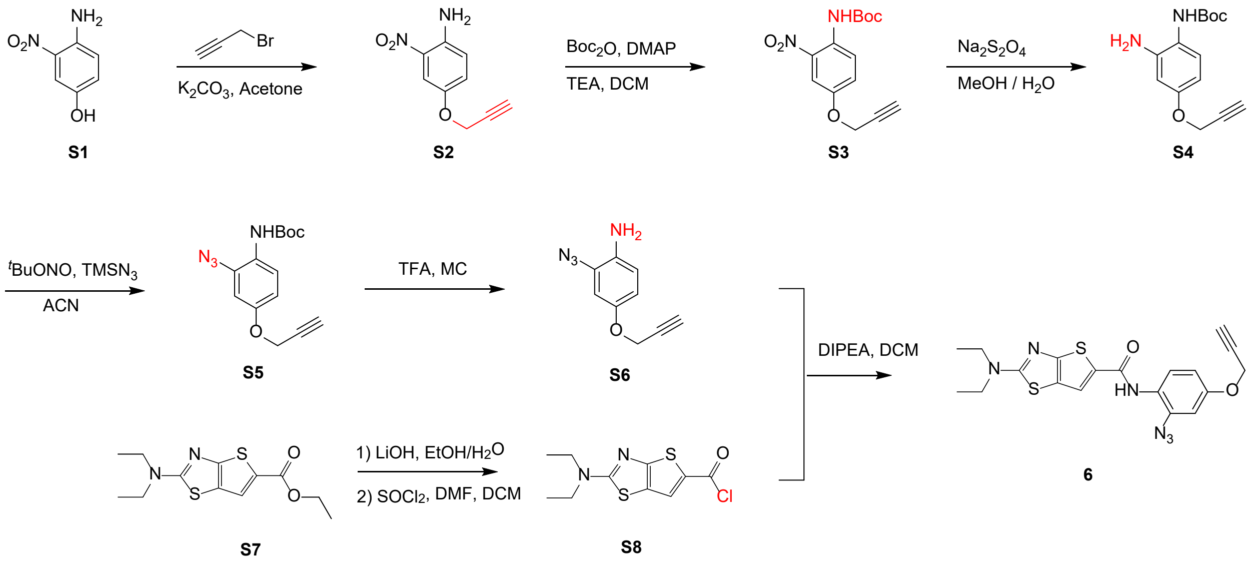

[Synthetic procedure]

*Synthesis of **S2***

To a solution of 4-amino-3-nitrophenol (**S1**, 1 g) in acetone (0.1 M) was added propargyl bromide (1.5 eq) and $K_2CO_3$ (2 eq). The reaction mixture was heated overnight at 60°C. After filtration of $K_2CO_3$, it was evaporated *in vacuo*. The crude mixture was purified by silica gel flash column chromatography (EA/Hex = 1/5 → 1/4) to yield **S2** as a yellow solid (85%). R*f* = 0.21 (EA/Hex = 1/4); $^{1}$H NMR (300 MHz, $CDCl_3$) δ 7.68 (d, *J* = 3.0 Hz, 1H), 7.13 (dd, *J* = 3.0 and 9.0 Hz, 1H), 6.81 (d, *J* = 9.0 Hz, 1H), 4.67 (d, *J* = 2.4 Hz, 2H), 2.55 (t, *J* = 2.4 Hz, 1H); $^{13}$C NMR (75 MHz, $CDCl_3$) δ 148.6, 140.6, 127.3, 120.3, 108.8, 78.0, 56.8 (t); LRMS (ESI) *m/z* 193 $[M+H]^+$.

*Synthesis of **S3***

To a solution of **S2** in DCM (0.2 M) was added Boc anhydride (1.1 eq), DMAP (0.2 eq), and TEA (1.2 eq). The reaction mixture was stirred overnight at room T. After completion of reaction, the reaction mixture was diluted with EA, washed with sat. $NH_4Cl$ and brine subsequently, dried over $MgSO_4$, and evaporated *in vacuo*. The crude mixture was purified by flash column chromatography (EA/Hex = 1/4→ 1/3) to yield **S3** as a yellow solid (40%). R*f* = 0.61 (EA/Hex = 1/4); $^1H$ NMR (300 MHz, $CDCl_3$) δ 9.42 (bs, 1H), 8.46 (d, *J* = 9.3 Hz, 1H), 7.76 (d, *J* = 3.0 Hz, 1H), 7.27 (dd, *J* = 3.0 and 9.3 Hz, 1H), 4.72 (d, *J* = 2.7 Hz, 2H), 2.55 (t, *J* = 2.7 Hz, 1H), 1.53 (s, 9H); $^{13}C$ NMR (75 MHz, $CDCl_3$) δ 152.5, 151.8, 136.4, 130.5, 124.5, 122.5, 110.3, 81.8, 76.6, 56.6, 28.3; LRMS (ESI) *m/z* 293 $[M+H]^+$.

*Synthesis of* ***S4***

To a solution of **S3** in MeOH (0.1 M) was added $Na_2S_2O_4$ (5 eq) solution in water (0.5 M). The reaction mixture was heated at 60°C for about 1 h until the yellow solution color disappeared. After evaporation the solvent in vacuo, it was resuspended in EA and washed with brine, dried over $MgSO_4$, and evaporated *in vacuo.* The crude mixture was purified by flash column chromatography (EA/Hex = 1/4 → 1/2 → 1/1) to yield **S4** as a pale-yellow liquid (42%). R*f* = 0.08 (EA/Hex = 1/4); $^1H$ NMR (300 MHz, $CD_3OD$) δ 6.97 (d, *J* = 8.7 Hz, 1H), 6.43 (d, *J* = 2.7 Hz, 1H), 6.31 (dd, *J* = 2.7 and 8.7 Hz, 1H), 4.62 (d, *J* = 2.7 Hz, 2H), 2.88 (t, *J* = 2.7 Hz, 1H), 1.49 (s, 9H); $^{13}C$ NMR (75 MHz, $CD_3OD$) δ 158.0, 157.0, 144.6, 128.4, 119.0, 105.7, 104.0, 80.8. 80.0, 76.5 (d), 56.6 (t), 49.9, 28.7; LRMS (ESI) *m/z* 263 $[M+H]^+$.

*Synthesis of* ***S5***

To a solution of **S4** in anhydrous ACN (0.2 M) was dropwise added t-butyl nitrite and azidotrimethylsilane at 0°C. The reaction mixture was stirred at room T for 1 h and then evaporated *in vacuo*. The crude mixture was purified by flash column chromatography (EA/Hex = 5/95 → 1/9) to afford a yellow solid (24%). R*f* = 0.56 (EA/Hex = 1/4); $^1H$ NMR (300 MHz, Acetone-*d*$_6$) δ 7.80 (d, *J* = 9.0 Hz, 1H), 7.23 (bs, 1H), 6.85–6.79 (m, 2H), 4.80 (d, *J* = 2.7 Hz, 2H), 3.09 (t, *J* = 2.7 Hz, 1H), 1.49 (s, 9H); $^{13}C$ NMR (75 MHz, Acetone-*d*$_6$) δ 155.1, 153.6, 131.3, 125.0, 123.5, 112.5, 106.3, 80.3, 79.5, 77.3, 77.2, 56.8 (t), 28.4.

*Synthesis of* ***6***

To a solution of **S5** in DCM (0.2 M) was added TFA (20% v/v). The reaction mixture was stirred at room T for about 2 h and then evaporated *in vacuo*. The reaction mixture was suspended in EA and sat. $NaHCO_3$

and the organic layer was separated. The organic layer was washed with brine, dried over $MgSO_4$, and evaporated *in vacuo* to yield the crude **S6** as a free base.

To a solution of ethyl 2-(diethylamino)thieno[2,3-*d*]thiazole-5-carboxylate (**S7**) in EtOH (0.3 M) was added aqueous LiOH solution (2 eq, 1 M). The reaction mixture was stirred overnight at room T. After evaporation of EtOH, the mixture was washed with EA to remove remaining starting material and organic impurities. The aqueous layer was adjusted pH with 1 N HCl down to 4–5. The desired product was extracted with EA three times and the combined organic layers were dried over $Na_2SO_4$ and evaporated *in vacuo* to afford a white solid. To suspension of this solid in anhydrous DCM (0.5 M) was added $SOCl_2$ (5 eq) and a few drops of anhydrous DMF. The reaction mixture was stirred for 1 h at room T and then evaporated completely to yield the crude **S8**.

To a solution of **S8** (1 eq) in DCM (1 M) was added **S6** solution in DCM (1 eq, 1 M) and DIPEA at 0°C. The reaction mixture was stirred at room T for 4 h. After completion of the reaction, the reaction mixture was diluted with EA, washed with brine, dried over $MgSO_4$, and evaporated *in vacuo*. The crude mixture was purified by flash column chromatography (EA/DCM = 5/95) and then recrystallized with acetone/$Et_2O$ to yield **6** as a pale-brown solid (63%). R*f* = 0.49 (EA/Hex = 1/1); $^1H$ NMR (300 MHz, $CDCl_3$) δ 8.30 (d, *J* = 9.6 Hz, 1H), 7.82 (s, 1H), 7.57 (s, 1H), 6.79–6.75 (m, 2H), 4.69 (d, *J* = 2.1 Hz, 2H), 3.54 (q, *J* = 7.1 Hz, 2H), 2.55 (t, *J* = 2.1 Hz, 1H), 1.28 (t, *J* = 7.1 Hz, 3H); $^{13}C$ NMR (75 MHz, $CDCl_3$) δ 173.4, 160.2, 159.8, 154.3, 129.8, 129.4, 124.1, 122.3, 122.2, 119.9, 111.4, 105.5, 78.3, 76.1, 56.4 (t), 45.9, 12.7; LRMS (ESI) *m/z* 427 $[M+H]^+$.

#### 4-4. Synthesis of radioactive $^{14}C$-labeled 4

[Synthetic scheme]

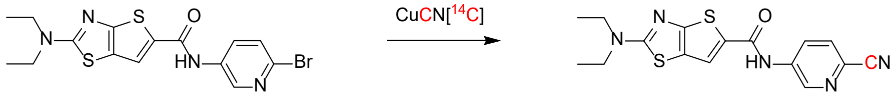

**S9** **4 [$^{14}C$]**

[Synthetic procedure]

The synthesis of radioactive DDU359[$^{14}C$] was optimized by normal CuCN first and then applied to the incorporation of real 14C-labeled CuCN[$^{14}C$] in radioactive room. Starting material **S9** was synthesized from commercially available ethyl 2-(diethylamino)thieno[2,3-*d*]thiazole-5-carboxylate (**S7**) and 3-amino-6-bromopyridine using the same procedure described above. To a solution of **S9** (25 mg) in DCM (5 mL) was added 0.2 mL of 33% HBr in AcOH. $Et_2O$ (10 mL) was added in the mixture and it was stirred for about 30 min at room T. The solid was filtered, washed with $Et_2O$, and dried under vacuum to yield HBr salt of **S7**. The dried HBr salt and CuCN[$^{14}C$] (56 mCi/mmol from American Radiolabeled Chemicals, Inc.) were suspended in 2 mL of anhydrous DMF in the heavy-wall pressure vessel, and then stirring overnight at 150°C. The color of reaction solution became dark greenish black. After checking the TLC to monitor the generation of desired product (TLC matched the dry-run pattern. R*f* = 0.34 of desired product and R*f* = 0.45 of remaining starting material at EA/Hex = 1/1), the reaction mixture was washed aq. $NH_4OH$ twice and the organic layer was dried under Ar gas. According to the results of several dry-run experiments, the purity was consistently confirmed ~80% by LC/MS and TLC. The final weight of product was 14.2 mg and it has approximately 2 mCi radioactivity. Due to radioactive material safety regulations, we used a crude mixture without quality control.

#### 4-5. Chemical oxidation test and confirmation of thiol adduct

[Synthetic scheme]

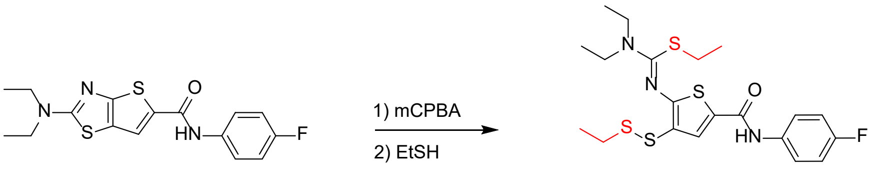

[Test procedure]

To a solution of **5** in MeOH/DCM (1/1, 0.2 M) was added mCPBA (2 eq). The solution was stirred for about 30 min at room T. The excess amount of EtSH (20 eq) was added to reaction mixture and stirred for another 30 min. The excess amount of EtSH was dried by air blowing. It was diluted with EA and washed with sat. $NaHCO_3$. The separated organic layer was dried over $NasSO_4$ and evaporated *in vacuo*. The crude mixture was dissolved in EtOH to make crystal **8** for x-ray crystallography. Yellow crystal with R*f* = 0.57 (EA/Hex = 1/2); $^1$H NMR (300 MHz, DMSO-*d*$_6$) $\delta$ 10.07 (s, 1H), 7.97 (s, 1H), 7.73–7.68 (m, 2H), 7.17 (t, *J* = 8.9 Hz, 2H), 3.65–3.61 (m, 4H), 2.79 (q, *J* = 7.2 Hz, 2H), 2.63 (q, *J* = 7.3 Hz, 2H), 1.30–1.12 (m, 12H); LRMS (ESI) *m/z* 472 $[M+H]^+$.

[NMR spectra]

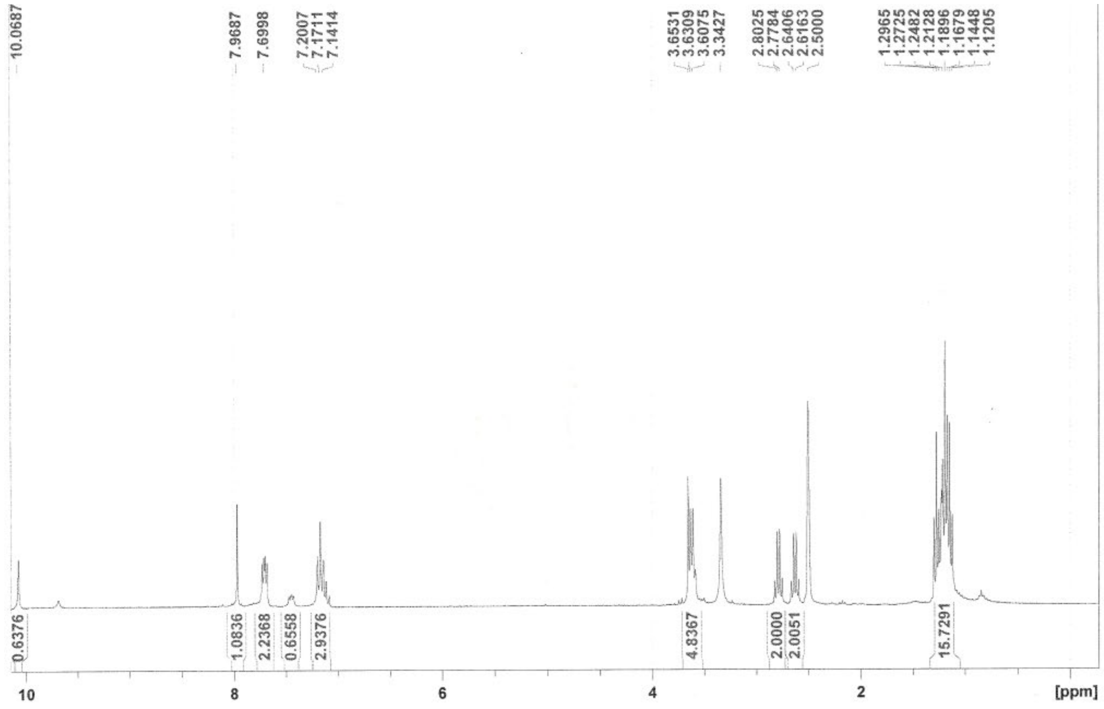
